# Agricultural digestate ultrafiltrate outperforms synthetic media for microalgal biomass production while enabling biostimulant generation

**DOI:** 10.64898/2026.10.29.762777

**Authors:** Tea Miotti, Mauro Maver, Marika Peli, Riccardo Lo Coco, Giacomo Rodegher, Giulia Viglioli, Nicola Frison, David Bolzonella, Anita Zamboni, Matteo Ballottari, Stefano Cazzaniga

**Author notes:** Corresponding author: Matteo Ballottari, Dipartimento di Biotecnologie, Università di Verona, Strada le Grazie 15, 37134 Verona Italy.

## Abstract

Livestock digestate ultrafiltrate (UF), obtained through membrane treatment of anaerobic digestate, is a nutrient-rich stream whose valorization remains largely unexplored. By removing suspended solids while retaining dissolved nutrients, ultrafiltration reduces turbidity and overcomes major limitations associated with using digestate for microalgal cultivation. This study evaluated UF as a nutrient source for *Chlorella sorokiniana* and *Nannochloropsis oceanica*, coupling ammonium recovery and biomass production with biomass valorization as a plant biostimulant. Given the high ammonium content of UF, ammonium tolerance was assessed in both species using ammonium bicarbonate as the nitrogen source. Increasing UF concentrations were then tested in laboratory photobioreactors and compared with synthetic media. Biomass productivity, nutrient removal, biomass composition, and biostimulant activity were evaluated. Both species tolerated ammonium-rich conditions and grew efficiently in UF-based media. UF significantly outperformed synthetic media, increasing biomass accumulation by up to 104% in *N. oceanica* and 133% in *C. sorokiniana*, while ammonium removal exceeded 99% and 98%, respectively. *C. sorokiniana* maintained high productivity when UF was diluted exclusively with water, indicating that UF supplied most nutrients required for growth. The enhanced productivity could not be explained solely by ammonium availability, suggesting a contribution from the complex nutrient composition retained during ultrafiltration. UF-derived biomass exhibited biostimulant activity, promoting tomato root development and increasing root and shoot biomass under reduced nitrogen availability. Digestate-derived UF therefore acts as both an efficient nutrient-recovery stream and a growth-promoting microalgal medium, enabling the conversion of membrane-treated digestate into biostimulant biomass within a circular biorefinery framework.

**Highlights:**

- Digestate ultrafiltrate outperformed synthetic media for algae cultivation
- *Chlorella sorokiniana* biomass increased by up to 133%
- *Nannochloropsis oceanica* biomass increased by up to 104%
- Ammonium removal reached >98% during microalgal cultivation
- Ultrafiltrate derived biomass promoted root and shoot development in tomato plants

## 1. Introduction

Anaerobic digestion has become one of the most widely adopted technologies for treating agricultural and livestock waste, generating renewable energy in the form of biogas (Sharma et al., 2025; Yan, 2025). Despite these advantages, the management of digestate remains a major challenge for the sustainability of anaerobic digestion systems (Bernhard et al., 2015; Chojnacka & Chojnacki, 2024; Herbes et al., 2020; Mancuso et al., 2024; Rizzioli et al., 2025). The liquid fraction of digestate contains substantial amounts of ammoniacal nitrogen, potassium, and other nutrients, making it a valuable resource but also posing environmental risks if improperly managed (Chojnacka & Chojnacki, 2024; Leong & Chang, 2022; Wester-Larsen et al., 2022). Excessive nutrient loading can promote ammonia volatilization, nitrate leaching and eutrophication of receiving water bodies, creating increasing pressure for the development of efficient nutrient recovery strategies.

Among the different approaches proposed for digestate valorization, microalgae-based technologies have attracted considerable attention because they combine wastewater treatment with the ability to efficiently consume nitrogen and phosphorus compounds, as well as CO2, thereby producing valuable biomass (Bauer et al., 2021; Fernandes et al., 2020; Fuentes-Grünewald et al., 2021; Gu et al., 2021; Leong & Chang, 2022; Manzo et al., 2026; Pizzera et al., 2019; Praveen et al., 2018; Ramasamy et al., 2024; Rossi et al., 2025; Sasongko et al., 2025; Scarponi et al., 2021; Sobolewska et al., 2023; Stiles et al., 2018; Tawfik et al., 2022; Uggetti et al., 2014; Xia & Murphy, 2016; Zhu et al., 2022). As photosynthetic microorganisms, not only do they contribute to carbon capture, but they also exploit light energy to produce the chemical energy in the form of ATP and NADPH required to efficiently assimilate inorganic nitrogen and phosphorus. The result is the conversion of wastes, and potentially pollutant compounds, into valuable biomass that can be exploited for multiple applications, including feed ingredients, biofuels, bioproducts, and agricultural biostimulants (Leong & Chang, 2022; Ljumovic et al., 2024; Miotti et al., 2026; Stiles et al., 2018; Tawfik et al., 2022; Xia & Murphy, 2016).

However, the direct cultivation of microalgae on anaerobic digestate is often limited by several physicochemical constraints. Raw digestate typically exhibits high turbidity, elevated solids content, microbial contamination and potentially inhibitory concentrations of ammoniacal nitrogen. These characteristics reduce light penetration, alter light distribution within the culture medium and may negatively affect microalgal productivity. Consequently, pretreatment steps are often required before digestate can be used efficiently as a cultivation substrate (Bauer et al., 2021; Chuka-ogwude et al., 2020; Fernandes et al., 2020; Miotti et al., 2026).

Membrane-based separation technologies have emerged as promising solutions to overcome these limitations. In particular, ultrafiltration removes suspended solids while retaining a large fraction of dissolved nutrients in the permeate, yielding a clarified ultrafiltrate with reduced turbidity and high concentrations of ammonium and potassium (Rizzioli et al., 2023; Rizzioli et al., 2025). This nutrient-rich stream is particularly attractive for microalgal cultivation because it simultaneously supplies essential macro- and micronutrients while improving light availability compared with untreated digestate (Fernandes et al., 2020; Veronesi et al., 2015). Nevertheless, studies focused specifically on the use of livestock digestate ultrafiltrates as complete nutrient sources for industrially relevant microalgae remain relatively scarce.

Despite the potential of digestate-derived streams, microalgal cultivation remains strongly species-dependent. High concentrations of ammoniacal nitrogen may represent both an opportunity and a challenge. Ammonium is generally considered the preferred nitrogen source for many microalgae because it can be directly incorporated into amino acids without the energetic cost of nitrate reduction (Sanz-Luque et al., 2015; Terrado et al., 2015). Nevertheless, excessive concentrations strongly influence the pH of the cultivation medium, potentially impairing photosynthesis, promoting oxidative stress, and inhibiting growth (Collos & Harrison, 2014; Freudenberg et al., 2021; Markou & Muylaert, 2016; Praveen et al., 2018). Therefore, identifying strains capable of efficiently utilizing nutrient-rich digestate fractions is a crucial prerequisite for process development.

Among the species currently investigated for wastewater valorization, *Chlorella* and *Nannochloropsis* are genera of particularly attractive candidates: different *Chlorella* species are recognized for their high growth rate, tolerance to stressful environmental conditions, and outstanding nutrient-removal capacity, making it one of the most promising microalgae for wastewater treatment applications (Alice et al., 2021; Dahmani et al., 2016; De-Bashan et al., 2008; Gu et al., 2021; Gupta et al., 2019; Li et al., 2011; Liang et al., 2010; Ljumovic et al., 2024). Species belonging to the *Nannochloropsis* genera are also industrially relevant: these are marine species characterized by high robustness and adaptability to saline environments, providing an interesting complementary platform for the exploitation of digestate-derived resources (Brooke et al., 2025; Hairuddin et al., 2025).

Beyond wastewater treatment and nutrient recovery, the valorization of the resulting microalgal biomass is critical to improve the economic viability of integrated digestate-based biorefineries. Microalgae synthesize a wide range of bioactive compounds, including amino acids, polysaccharides, phytohormones, vitamins, pigments and antioxidant molecules that can positively influence plant growth and stress tolerance (Martini et al., 2021; Mutale-Joan et al., 2020; Álvarez-González et al., 2025). Consequently, microalgal biomass has recently attracted increasing interest as a sustainable source of agricultural biostimulants capable of enhancing nutrient uptake, root development and crop productivity while reducing reliance on synthetic agrochemicals. Numerous studies have demonstrated positive effects of microalgae-derived products on seed germination, root architecture and plant performance under both optimal and stress conditions (Chiaiese et al., 2018; Kapoore et al., 2021; Martini et al., 2021; Morillas-España et al., 2022; Mutale-Joan et al., 2020).

The integration of microalgal cultivation with digestate treatment offers a particularly attractive circular approach. Nutrients recovered from agricultural wastes can be converted into algal biomass and subsequently reintroduced into crop production systems as biostimulant products, thereby closing nutrient loops and increasing the value generated from waste streams. However, most studies have focused on nutrient removal and biomass production using raw or diluted digestates, while nutrient-rich streams from membrane-based treatments remain comparatively understudied. Furthermore, the potential to couple digestate-derived nutrient recovery with the production of microalgal biomass for biostimulant applications has received little attention. Consequently, the development of integrated circular biorefinery systems that maximize both environmental and economic benefits remains largely unexplored.

In this study, we evaluated the potential of an ultrafiltrate derived from livestock anaerobic digestate as an alternative nutrient source for cultivating the microalgal species *Chlorella sorokiniana* and *Nannochloropsis oceanica*. *C. sorokiniana* has been widely documented for wastewater treatment and high growth rate for biomass production (De-Bashan et al., 2008). *N. oceanica*, a marine microalga, offers advantages through its ability to cultivate in saline media and under low-light conditions (Huete-Ortega et al., 2018). The ammonium tolerance of *Chlorella sorokiniana* and *Nannochloropsis oceanica* was first assessed to identify suitable operating conditions for the utilization of ultrafiltrate. Subsequently, the effect of increasing ultrafiltrate concentrations on biomass productivity and nutrient removal was investigated, and the ability of ultrafiltrate to replace conventional synthetic media was assessed. Finally, a valorization pathway for the biomass produced was investigated by successfully exploring its potential biostimulatory activity on crop plants. The results demonstrate that digestate-derived ultrafiltrate can simultaneously support efficient ammonium recovery and enhanced biomass production, providing a promising route for integrating wastewater treatment and microalgal biorefineries within circular bioeconomy frameworks.

## 2. Materials and methods

### 2.1 Digestate ultrafiltrate

The nutrient solution used for microalgal cultivation was obtained from the liquid fraction of anaerobic digestate collected at a full-scale anaerobic digestion plant located in Northern Italy (Isola della Scala, Verona, Italy), where bovine manure and crop residues are co-digested under mesophilic conditions (around 40°C). The digestate was first screened through a 500 µm mesh to remove coarse suspended solids. It was then ultrafiltered through ceramic membranes with a molecular weight cut-off (MWCO) of 150 kDa, yielding a clarified, nutrient-rich ultrafiltrate (UF). The main physicochemical characteristics of the UF are reported in Table 1

**Table 1.** Analysis of ultrafiltrate composition.

|  | pH | COD<br>mgCOD<br>l<sup>-1</sup> | TKN<br>mgN l<sup>-1</sup> | NH<sub>4</sub>-N<br>mgN l<sup>-1</sup> | Fe<br>mg L<sup>-1</sup> | B<br>mg L<sup>-1</sup> | Mg<br>mg L<sup>-1</sup> | P<br>mg L<sup>-1</sup> | K<br>mg<br>L<sup>-1</sup> | Mn<br>mg<br>L<sup>-1</sup> | Ni<br>mg<br>L<sup>-1</sup> | Cu<br>mg<br>L<sup>-1</sup> | Zn<br>mg<br>L<sup>-1</sup> |
| --- | --- | --- | --- | --- | --- | --- | --- | --- | --- | --- | --- | --- | --- |
| Ultrafiltrate<br>(UF) | 7.5 | 8256±212 | 4012±92 | 3753±112 | 3.66±0.08 | 2.56 ± 0.01 | n.d | 87.92 ± 0.67 | 4886.94 ± 135.35 | trace | 0.40 ± 0 | trace | trace |

Depending on the experimental conditions, the UF was diluted to the desired concentrations with either modified synthetic culture medium or sterile water. No sterilization treatment was applied to the UF prior to the experiments. To assess nutrient removal efficiency during microalgal growth, Total Kjeldahl Nitrogen (TKN), ammoniacal nitrogen (NH4-N), and Chemical Oxygen Demand (COD) were determined at the beginning and at the end of the cultivation period (Cavinato et al., 2010; Giuliano et al., 2013). The concentrations of phosphorus, potassium, magnesium, calcium and trace metals were quantified by inductively coupled plasma mass spectrometry (ICP-MS) on three independent replicates (n = 3).

### 2.2 Microalgal strains and culture conditions

*C. sorokiniana* (CS) was obtained from UTEX colture collection (UTEX1230) while *N. oceanica IMET1* was gifted from Prof. Sarah D’Adamo, Wageningen University and Research (WUR). For all experiments, *C. sorokiniana and N.oceanica* were grown in liquid synthetic minimum (BG-11) medium (Allen & Stanier, 1968) and artificial seawater (ASW) (Cai et al., 2013) respectively until reaching the required inoculum concentration for each experiment. Cell cultures were maintained at 25 °C under a light intensity of 120 μmol m^−2^ s^−1^ and a photoperiod of 16 h light / 8 h dark. No additional inorganic carbon was supplied, and a rotative shaker (150 rpm) was used to prevent cell sedimentation and induce gas exchange.

All cultivations were conducted in batch mode in 80 mL airlift photobioreactors (PBRs) in the Multicultivator system MC-1000-OD (PSI, Czech Republic) with independent white light-emitting diodes (LEDs) illumination, temperature control at 24 °C and continous 3% CO_2_ insufflation. Cultures were initially exposed to 100 μmol photons m⁻² s⁻¹ for 12 h, followed by a gradual increase in light intensity to 1000 μmol photons m⁻² s⁻¹ over the subsequent 24 h. Cultures were then maintained at 1000 μmol photons m⁻² s⁻¹ for the remaining cultivation period, until the saturation phase was reached. Cell density was measured every two days with a cell counter (Multisizer 4e, Beckman Coulter).

In order to test tolerance to ammonium concentration, *C. sorokiniana* was cultivated in modified BG-11 medium at nitrogen concentrations of 0.24, 0.48, 0.98 and 1.48 g N L⁻¹ in three independent biological replicates for each cultivation condition. Similarly, *N. oceanica* was cultivated in modified ASW medium at nitrogen concentrations of 0.06, 0.18, 0.72, 1.08 and 1.44 g N L⁻¹. In both cases, sodium nitrate (NaNO₃) was omitted from the synthetic media to ensure that ammonium bicarbonate was the supplied nitrogen source. Cultures grown in complete BG-11 and ASW media, with nitrate as N source, were used as controls for *C. sorokiniana* and *N. oceanica*, respectively. All experiments were performed under the cultivation conditions previusly described. Growth was monitored by measuring OD₇₂₀ and cell concentration, while final biomass accumulation was determined as dry weight (DW) on the last day of growth.

The ability of *C. sorokiniana* and *N. oceanica* to utilize UF as an alternative nutrient source was investigated by cultivating both species at increasing UF concentrations. The dilution ranges were selected based on each strain’s ammonium tolerance, as determined in preliminary experiments. *N. oceanica* was cultivated in modified ASW medium prepared without sodium nitrate but supplemented with 5, 10, 15 and 20% (v/v) UF. Cultures grown in complete ASW medium without UF were used as controls. Cultivation was performed for 17 days, after which final biomass concentration was determined as DW. Similarly, *C. sorokiniana* was cultivated in modified BG-11 medium prepared without sodium nitrate but supplemented with 15, 20, 30 and 40% (v/v) UF, using complete BG-11 medium as the control. To evaluate if UF could provide the nutrients required to sustain microalgal growth without supplementation with conventional synthetic culture media, UF was supplied at 20% and 30 % for *N. oceanica* and *C. sorokiniana*, respectively, and diluted in sterilized water.

To determine the biomass dry weight (DW), at the end of the growth, a 50 mL culture sample was collected and centrifuged (3300 xg, 10 m, RT), the supernatant was discarded, the pellet sample was washed with water in order to remove residual salts, lyophilized for 48 h and then weighted. Maximal volumetric biomass productivities (P_max_) at the cultivation time t were calculated upon fitting of growth curves (OD_720_) by the sigmoidal Boltzman model function. The fitted curves were then used to calculate the 1^st^ derivative function. The median and the maximum value of the 1^st^ derivative function was used to determine the maximum biomass productivity from the average biomass productivity as previously reported (Canini et al., 2024).

### 2.3 Evaluation of biostimulant activity of microalgae

#### 2.3.1 Chemical characterization of microalgae

Total nitrogen (N) and carbon (C) contents were determined by CHN analysis using 100 mg of lyophilized biomass (Table S1). Macro- and micronutrient concentrations (Ca, K, Mg, P, Cu, Fe, Mn, Mo, Ni, and Zn, Table S2) were measured in mineralized samples obtained by microwave-assisted acid digestion. Approximately 20 mg of each sample was weighed into a TFM micro-sampling insert (Milestone Srl, Bergamo, Italy) and digested with 350 µL of 60% ultrapure HNO□ (Romil Ltd, Cambridge, UK). The inserts were placed inside 100-mL digestion vessels containing 10 mL of ultrapure water and 1 mL of 30% ultrapure H□O□ (Romil Ltd, Cambridge, UK). As a certified reference material, approximately 20 mg of NIST 1515 (apple leaves) was digested alongside the samples. Digestion was performed using a microwave digestion system (Start D, Milestone Srl, Bergamo, Italy). Samples were heated to 180 °C over a 20-min ramp period and maintained at this temperature for a further 20 min. After digestion, the solutions were diluted with ultrapure water to a final HNO□ concentration of 1%. Multi-element analysis was conducted using an iCAP™ RQ ICP-MS instrument (Thermo Fisher Scientific, Waltham, MA, USA). Macro- and micronutrient concentrations were quantified using multi-element calibration standards (Romil Ltd, Cambridge, UK). Both analyses were carried out for biomass of *Nannochloropsis oceanica* and *Chlorella sorokiniana* grown on the synthetic minimum medium (BG-11) (NO and CS) and on ultrafiltrate (NOUF and CSUF).

#### 2.3.2 Seed priming experiment

Tomato seeds (*Solanum lycopersicum* cv. Roma, Sementi Franchi) were surface-sterilized by immersion in a 5% (v/v) NaClO solution for 5 min, followed by three consecutive washes with sterile Milli-Q water (10 min each). The seeds were then air-dried under a laminar flow hood for 1 h. In parallel, algal suspensions were prepared by resuspending algal biomass in sterile water to different concentrations (0.02, 0.1, 1, 10, and 25 g L^-1^). The experimental design included three treatment groups: (i) control, in which seeds were not exposed to any priming solution; (ii) a control (0 g L^-1^), consisting of sterile water only; and (iii) algal biomass treatments at the selected concentrations. Surface-sterilized seeds were immersed in 50 mL of the respective priming solution at room temperature in the dark, with continuous agitation, for 24 h. Following priming, the solutions were discarded, and the seeds were dried under a laminar flow hood for a further 24 h before sowing. For each treatment, four Petri dishes containing 20 seeds each were prepared per experimental replicate. Seeds were placed on sterile filter paper moistened with 1 mL of sterile Milli-Q water and incubated in the dark at 23 °C for 4 days. At the end of the incubation period, germination percentage (GP) was calculated as the proportion of germinated seeds relative to the total number of seeds, expressed as a percentage. Radicle length of germinated seeds was measured using WinRHIZO^TM^ image analysis software. The experiment was repeated independently 3 times, with 4 plates per treatment. For each replicated experiment, the mean values of total seedlings for four plates were calculated for root length and germination %.

#### 2.3.3 Treatment of tomato seedlings in hydroponics

To evaluate the biostimulant effect of algal biomasses, hydroponic trials were conducted on tomato plants (*Solanum lycopersicum* L. var. Roma). Tomato seeds were surface-sterilized by immersion in a 5% NaClO solution for 5 min, followed by three successive 10-min washes with sterile Milli-Q water. Seeds were then placed in sterile water at 4°C for 48 h to synchronise germination. Following this step, the sterilised seeds were arranged in Petri dishes containing plant agar (8 g/L) and incubated at 23°C. After 5 days, 40 seedlings per treatment (20 per tray) were transferred to trays containing 550 mL of aerated nutrient solution, continuously oxygenated by air insufflation, and the experimental treatments were applied. The nutrient solution was composed of the following macronutrients: 0.7 mM K_2_SO_4_, 0.5 mM MgSO_4_, 0.1 mM KCl, 0.1 mM KH_2_PO_4_, 2 mM Ca(NO_3_)_2_, 50 μM Fe-EDTA, 0.5 μM MnSO_4_, 0.5 μM ZnSO_4_, 0.2 μM CuSO_4_, 10 μM H_3_BO_3_, and 0.01 μM (NH_4_)_6_Mo_7_O_24_ (pH 6.0). The experimental design included two control treatments: an untreated control containing only nutrient solution, and a positive control consisting of a commercial formulation based on *Ascophyllum nodosum*, normalised to supply the same amount of carbon (C) as the microalgal biomasses. Algal biomasses were resuspended in water at a 1:10 (w/v) ratio and added to the nutrient solution (drench application) at a concentration of 47 mg C/L, corresponding to approximately 100 mg/L of algal biomass. After 7 days of growth under treatment, plants were harvested and analysed for root system length and surface area, lateral root length, and fresh biomass (FW) of both the root and shoot systems using WinRHIZO^TM^ software. The experiment was independently repeated two times (i.e., two separate growth and treatment cycles).

#### 2.3.4 Treatment of tomato seedlings in hydroponics carried out with different N availability

Tomato seeds (*Solanum lycopersicum* cv. Roma, Sementi Franchi) were germinated on paper towels soaked in 0.5 mM CaSO_4_ and kept in the dark at 22–26 °C for 5 days. Six seedlings of uniform size were then transferred to 2-L pots containing 1.8 L of aerated nutrient solution. Two nutritional regimes were tested: a full-strength nutrient solution (control) and a solution with 30% of the standard nitrogen content (30% N); each regime was tested with and without the addition of NUF, for a total of four treatments; in the NUF treatments, 100 mg of NUF was added to the nutrient solution at each renewal (55.56 mg L^-1^). The full nutrient solution contained the following macro- and micronutrients: 0.7 mM K_2_SO_4_, 0.5 mM MgSO_4_, 0.1 mM KCl, 0.1 mM KH_2_PO_4_, 2 mM Ca(NO_3_)_2_, 50 μM Fe-EDTA, 0.5 μM MnSO_4_, 0.5 μM ZnSO_4_, 0.2 μM CuSO_4_, 10 μM H_3_BO_3_, and 0.01 μM (NH_4_)_6_Mo_7_O_24_ (pH 6.0). In the two control treatments (100% N and 30% N), the nitrogen content was adjusted by supplementing the medium with Ca(NO₃)₂ to match the amount of nitrogen supplied to the corresponding NUF-treated cultures. The solution was continuously aerated and renewed every 6 days. Plants were grown in a growth chamber under a photosynthetic photon flux density (PPFD) of 200 μmol m^−2^ s^−1^, a 16 h/8 h day/night photoperiod, day/night temperatures of 25/22 °C, and 70% relative humidity, and were harvested 17 days after transfer to the nutrient solution. The experiment was independently repeated three times (i.e., three separate growth and treatment cycles). At harvest, each plant was separated into shoot and root fractions, and FW and dry weight (DW) was recorded for both; DW was subsequently determined after oven-drying at 65 °C for 3 days. Before drying, roots were scanned and analysed for the main root architectural parameters using WinRHIZO™ software (EPSON V850 Pro scanner; WinRHIZO™ Pro2021a; Regent Instruments Inc., Quebec, Canada). The root-to-shoot ratio was calculated for each treatment from the fresh and dry biomass values.

### 2.4 Statistics analysis

All the experiments described were performed on at least three independent biological replicates. The specific number of independent biological replicates is reported for each experiments. Statistical analysis was performed by using a two-sided Student’s *t*-test or one-way ANOVA with post-hoc Tukey test in the case of multiple comparisons. Error bars indicate standard deviation (SD). Significance thresholds were set to 0.05.

## 3. Results and discussion

### 3.1 Cultivation of Chlorella sorokiniana and Nannochloropsis oceanica in ultrafiltrate-based media

To evaluate the optimal condition for microalgal growth in ammonia-rich ultrafiltrate (UF), two microalgal species, *Chlorella sorokiniana* and *Nannochloropsis oceanica*, were selected based on their high tolerance to stressful conditions and their capacity to grow in high-ammonia environments (Collos & Harrison, 2014; Huete-Ortega et al., 2018).

Preliminary cultivation experiments were performed using ammonium bicarbonate (NH₄HCO₃), the same ammonium source present in the UF, to determine the tolerance limits of the two microalgal species to ammonia while excluding possible limitations or inhibitory effects associated with the UF matrix. Both microalgal species exhibited high tolerance to ammoniacal nitrogen, maintaining active growth at elevated NH₄HCO₃ concentrations (Fig. S1). *C. sorokiniana* was able to grow at ammonium nitrogen concentrations up to 0.98 g L⁻¹, corresponding to approximately one third of the ammonium present in the UF. The highest biomass production was obtained at 0.48 g L⁻¹ NH₄HCO₃, reaching a final dry weight of approximately 4.3 g L⁻¹, the 0.23 g L⁻¹ treatment also supported efficient growth, while concentrations above 0.98 g L⁻¹ resulted in complete growth inhibition, suggesting that excessive ammonium availability hamper *C. sorokiniana* cultivation.

Similarly, *N. oceanica* exhibited a high tolerance to ammonium supplementation, sustaining growth up to 0.72 g L⁻¹ (Fig. S1). Among the tested conditions, 0.18 g L⁻¹ NH₄HCO₃ resulted in the highest growth performance, with a final dry biomass concentration of 4.82 g L ⁻¹ (Fig. S1). Interestingly, cultures supplemented with 0.72 g L⁻¹ nitrogen exhibited a delayed transition to the stationary phase, reaching it approximately three days later than cultures with lower ammonia concentrations, indicating that increased nitrogen availability prolonged the active growth period.

Based on the concentration of ammonium nitrogen, supplied as NH□ HCO□, different ultrafiltrate (UF) dilutions, from 5 to 20%, were tested to evaluate their effect on *N. oceanica* cultivation in lab-scale photobioreactors. UF was diluted in modified ASW media without NaNO□ in order to exploit the ammonium in the UF as the sole nitrogen source. Cells grown in complete ASW medium were used as controls. Cultures were maintained at 25 °C under continuous aeration enriched with 3% CO₂. Light intensity was initially set at 100 μmol photons m⁻² s⁻¹ for 12 h and then gradually increased to 1000 μmol photons m⁻² s⁻¹ over the following 24 h. Cultures were subsequently maintained under these conditions until reaching the stationary phase. Growth was monitored by measuring both optical density at 720 nm (OD₇₂₀) and cell density (Fig. 1A, B). Cultivation in UF-based media markedly enhanced biomass accumulation compared with the synthetic control. During the early stages of growth, all UF treatments exhibited a lag phase that became more pronounced as UF concentration increased. However, after this initial adaptation period, all UF-supplemented cultures reached higher biomass levels than the control. Consistent with OD measurements, cultures grown with 10-20% UF displayed higher cell densities and biomass accumulation than the control. After 18 days of cultivation, the control reached a biomass concentration of approximately 3 g L⁻¹, whereas UF-treated cultures achieved 7-8 g L⁻¹, corresponding to more than a twofold increase in biomass yield. The highest productivity (approximately 2.2 g L⁻¹ d⁻¹) was observed in cultures supplemented with 10-15% UF, whereas the control did not exceed 1.0 g L⁻¹ d⁻¹. Although cultures grown with 20% UF ultimately reached similarly high biomass concentrations, their longer lag phase and slower growth kinetics resulted in lower overall productivity. Nevertheless, biomass productivity remained comparable to that obtained in the synthetic medium. Overall, these results demonstrate that *N. oceanica* can efficiently utilize digestate-derived UF as a nutrient source and that UF supplementation significantly enhances both biomass productivity and final biomass accumulation compared with conventional synthetic medium. None of the tested UF concentrations resulted in growth limitations. Conversely, the addition of UF promoted biomass accumulation, with all treatments showing higher final biomass production than the control. The highest increase in dry weight was observed at 10% and 15% UF supplementation, reaching up to a 104% improvement compared with the control (Fig. 1C). This enhanced biomass production was also reflected in the growth performance, with the 10% and 15% UF treatment showing the highest maximum productivity (Fig. 1D). 20% ultrafiltrate, on the other hand, positively affected biomass production, but the high ammonia concentration and low light penetration increased the lag phase, reducing maximum productivity (Fig. 1D) compared with lower UF conditions.

**Figure 1.**
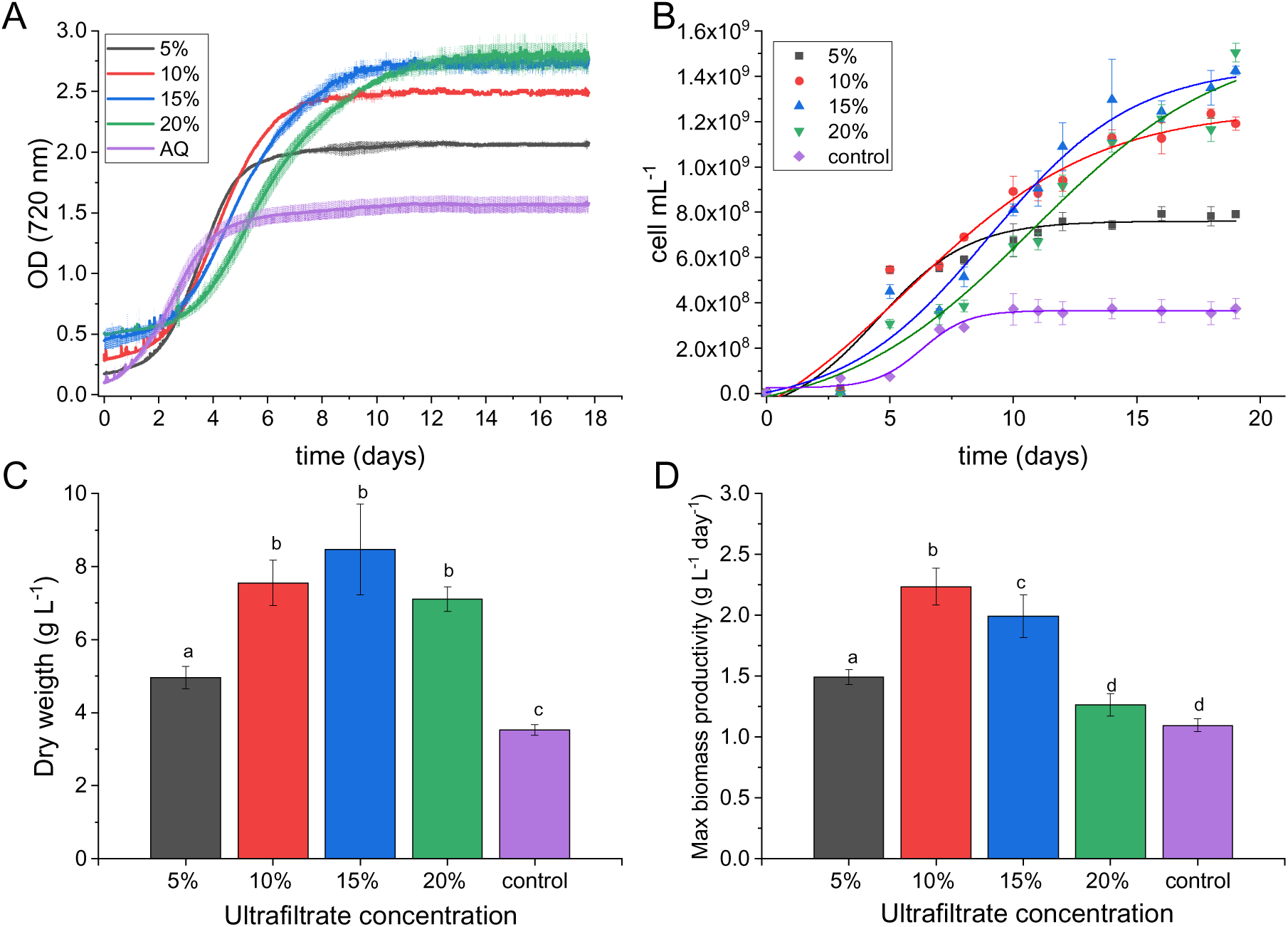
Growth performance of *Nannochloropsis oceanica* in digestate-derived ultrafiltrate (UF)-based media. Growth of *N. oceanica* cultivated in artificial seawater (ASW) supplemented with increasing concentrations of UF. The control culture (0% UF) was grown in complete ASW medium. Growth was monitored by measuring optical density at 720 nm (A) and cell density (B). Final biomass concentration was determined as dry weight at the end of the cultivation period (C), and maximum biomass productivity was calculated from growth kinetics and final dry weight values (D). The higher initial OD₇₂₀ observed in UF-containing cultures reflects the intrinsic turbidity and coloration of the ultrafiltrate rather than algal biomass accumulation. Data are presented as mean ± SD (n = 4). Different letters indicate statistically significant differences among treatments within each panel, according to one-way ANOVA followed by Tukey’s post-hoc test (p < 0.05).

Additional experiments were performed to determine whether UF alone could provide all the necessary components to sustain microalgal growth. Three UF-based conditions were compared with the ASW reference medium: 20% UF diluted with ASW, 20% UF diluted with tap water, and 20% UF diluted with tap water supplemented only with the NaCl needed for the marine strain (Fig. 2). As shown by OD₇₂₀ measurements and cell density (Fig. S2), cultures grown in 20% UF supplemented with ASW showed continuous growth, reaching a final biomass concentration of approximately 6.7 g L⁻¹ dry weight (Fig. S2). In contrast, both treatments containing UF diluted with water, either with or without NaCl addition did not grow. The inability of NaCl supplementation alone to restore growth suggests that growth limitation was not exclusively associated with osmotic conditions or salinity. To identify the nutrients limiting algal growth, individual salts were sequentially omitted from the ASW medium used for the dilution of UF, specifically removing Na_2_SO_4_, K_2_SO_4_, MgCl_2_ or CaCl_2_ (Fig. S3). The results clearly identified MgCl₂ and CaCl₂ as essential components in ASW salts for sustaining *N. oceanica* growth which cannot be properly substituted with UF. Consistent with the compositional analysis of the ultrafiltrate, which revealed only trace amounts of magnesium and calcium, these findings suggest that UF alone cannot provide sufficient concentrations of these elements to meet the physiological demands of the species.

**Figure 2.**
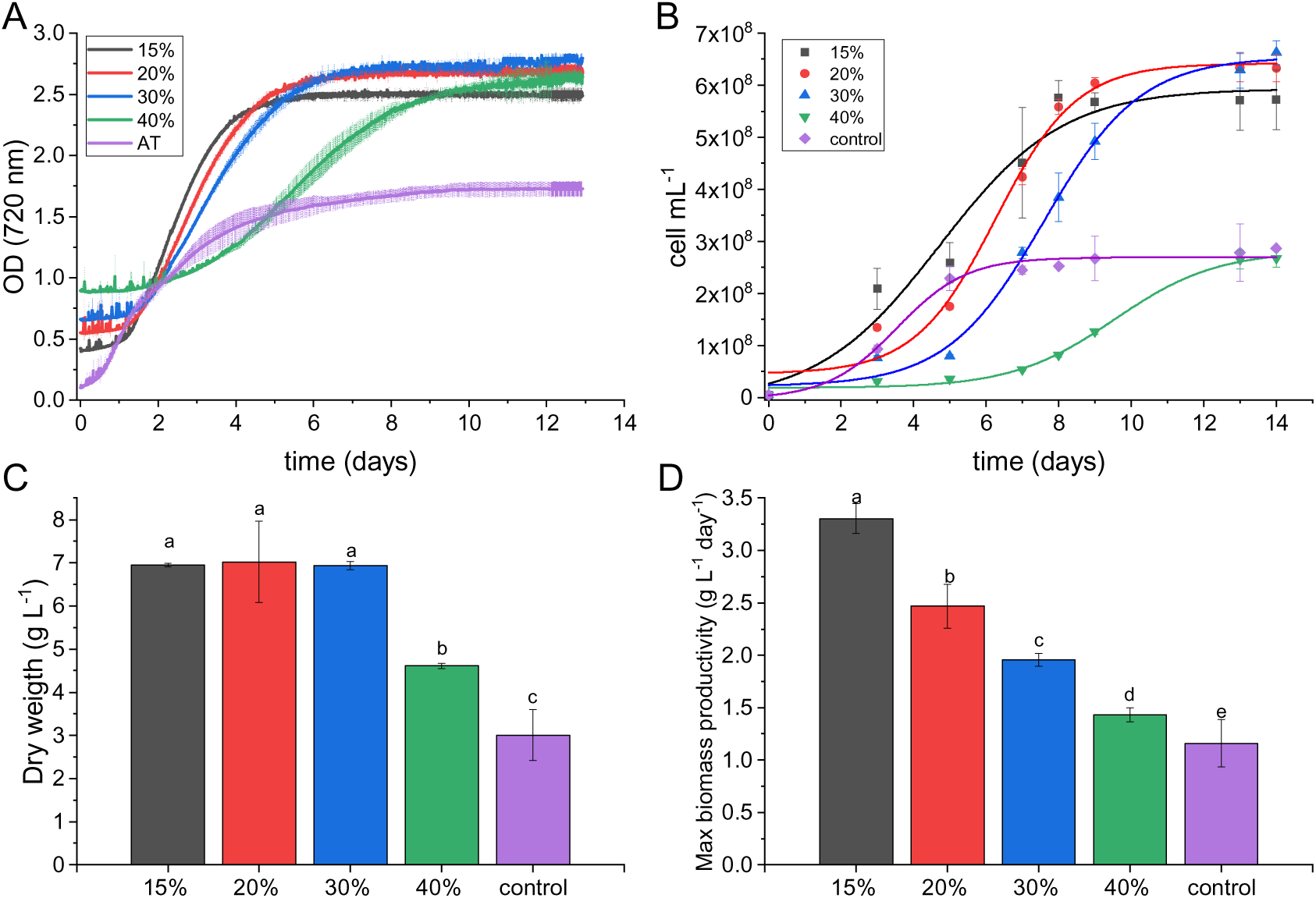
Growth performance of *Chlorella sorokiniana* in digestate-derived ultrafiltrate (UF)-based media. Growth of *C. sorokiniana* cultivated in BG-11 medium supplemented with increasing concentrations of UF. The control culture (0% UF) was grown in complete BG-11 medium without UF supplementation. Growth was monitored by measuring optical density at 720 nm (A) and cell density (B). Final biomass concentration was determined as dry weight at the end of the cultivation period (C), and maximum biomass productivity was calculated from growth kinetics and final dry weight values (D). The higher initial OD₇₂₀ observed in UF-containing cultures reflects the intrinsic turbidity and coloration of the ultrafiltrate rather than algal biomass accumulation. Data are presented as mean ± SD (n = 4). Different letters indicate statistically significant differences among treatments within each panel, according to one-way ANOVA followed by Tukey’s post-hoc test (p < 0.05).

The potential of UF as a cultivation substrate was also evaluated in *C. sorokiniana*. Based on the ammonium tolerance experiments, cultures were supplemented with UF as the sole N source at concentrations ranging from 15% to 40%, corresponding to total nitrogen concentrations between 0.49 and 1.48 g L⁻¹. Notably, ammonium concentrations above 0.98 g L⁻¹ were previously found to strongly inhibit the growth of *C. sorokiniana* (Fig. S1). Growth performance was compared with that obtained in the synthetic BG-11 medium. Similar to the observations for *N. oceanica*, cultures grown in BG-11 exhibited a rapid increase in biomass from the first day of cultivation, whereas UF-supplemented cultures displayed an initial lag phase. However, following this adaptation period, all cultures containing 15-30% UF outperformed the control, reaching substantially higher cell densities and biomass concentrations (Fig. 2). In particular, cell density increased to approximately 6 × 10□ cells mL⁻¹ in the 15-30% UF treatments, compared with 2.5 × 10□ cells mL⁻¹ in the BG-11 control. Consistently, final biomass concentration increased from approximately 3 g L⁻¹ in the control to about 7 g L⁻¹ in UF-grown cultures, corresponding to more than a twofold increase in biomass production. The highest UF concentration tested (40%) also supported greater biomass accumulation than the control, reaching approximately 4 g L⁻¹, although growth was lower than that observed in cultures supplemented with 15-30% UF. Biomass productivity followed a similar trend, with all treatments containing 15-30% UF outperforming the synthetic medium. The highest productivity (3.2 g L⁻¹ d⁻¹) was achieved with 15% UF, while increasing UF concentrations progressively reduced productivity due to slower growth kinetics. Nevertheless, even at 40% UF, biomass productivity remained comparable to that obtained in the BG-11 control (Fig. 2D). These findings indicate that the detrimental effects associated with elevated ammonium concentrations were largely compensated by the beneficial properties of UF. While the substitution of nitrate with ammonium in a synthetic medium improved biomass accumulation only at low nitrogen concentrations (0.48 g L⁻¹; Fig. S1), cultures supplemented with UF achieved substantially higher biomass yields even at comparable or higher nitrogen levels. Therefore, the growth-promoting effect of UF is unlikely to be attributable solely to ammonium availability and is more likely attributable to additional nutrients and growth-supporting compounds naturally present in the digestate-derived ultrafiltrate.

We further evaluated the ability of *C. sorokiniana* to grow on UF without synthetic nutrients (Fig. 3A-C). Cultures were grown either in 30% UF supplemented with BG-11 salts (without nitrate) or in 30% UF diluted exclusively with tap water. In contrast to *N. oceanica*, *C. sorokiniana* maintained robust growth when UF was provided as the sole nutrient source. The two treatments exhibited virtually identical growth kinetics and reached comparable final biomass concentrations of approximately 10 g L⁻¹ dry weight (Fig. 3C). These results indicate that the ultrafiltrate contains sufficient amounts of the essential macro- and micronutrients required to sustain *C. sorokiniana* growth, highlighting its potential as a cost-effective alternative to conventional synthetic media. Moreover, no statistically significant differences in final biomass production were observed among cultures supplemented with 15-30% UF, all of which achieved biomass yields up to 133% higher than those obtained in the synthetic control medium. Overall, these results demonstrate that digestate-derived UF can effectively replace conventional synthetic nutrients for the cultivation of *C. sorokiniana*, significantly enhancing both biomass accumulation and productivity over a broad range of ultrafiltrate concentrations. Differently, from N. oceanica the low concentrations of Mg^2+^ and Ca^2+^ in UF, combined with the presence of these elements in tap water, is sufficient to support the growth of *C. sorokiniana*.

**Figure 3.**
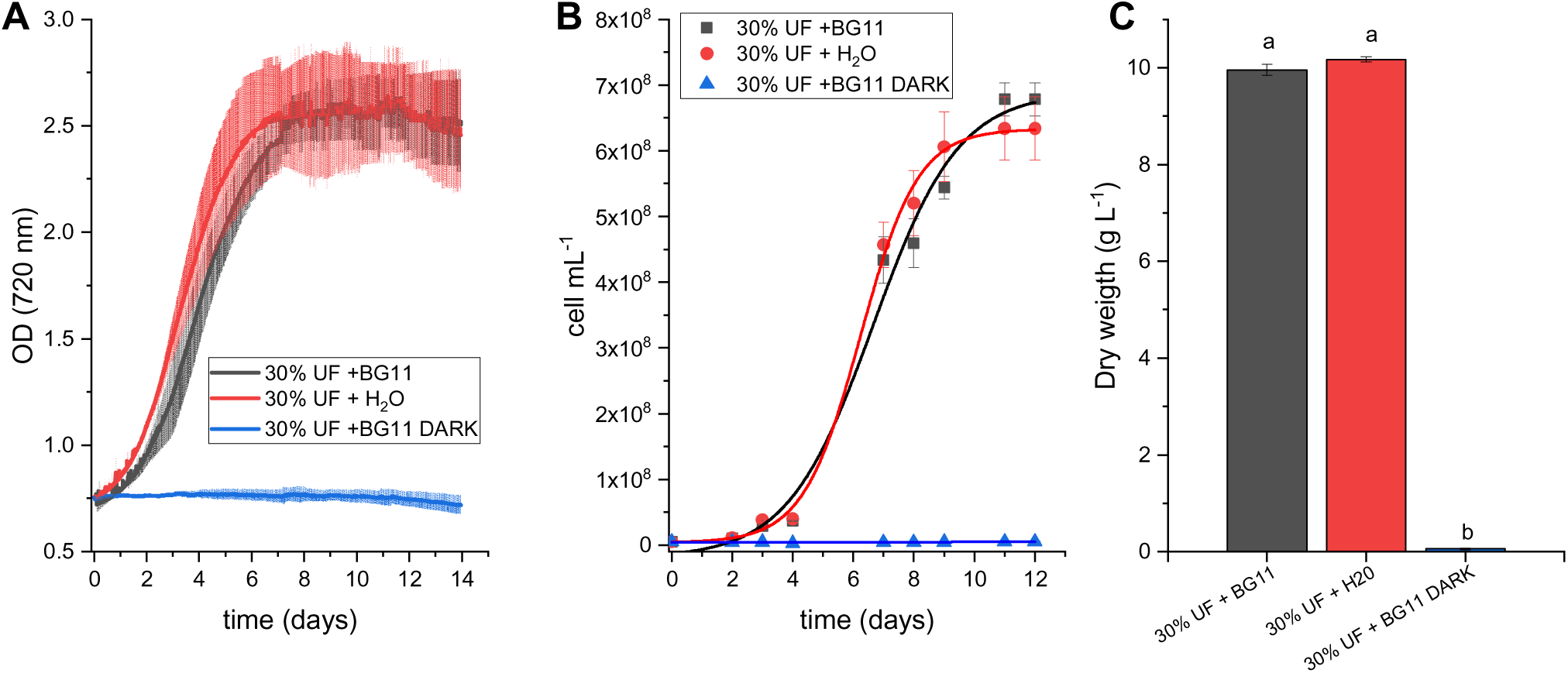
Growth of *Chlorella sorokiniana* in ultrafiltrate-based media without additional nutrient supplementation. Growth of *C. sorokiniana* cultivated in 30% ultrafiltrate (UF) supplemented with BG-11 salts (30% UF + BG-11), diluted exclusively with water (30% UF + H₂O), or grown in the dark in UF-supplemented BG-11 medium (30% UF + BG-11 Dark). Growth was monitored by measuring optical density at 720 nm (A) and cell density (B). Final biomass concentration was determined as dry weight at the end of the cultivation period (C). Data are presented as mean ± SD (n = 2).

As *C. sorokiniana* is a facultative mixotrophic microalga capable of utilizing organic carbon sources, an additional experiment was performed to assess whether the enhanced biomass accumulation observed in UF cultures could be attributed to the assimilation of residual organic compounds present in the ultrafiltrate (Table 1). To this end, cells were cultivated in 30% UF supplemented with BG-11 salts under complete darkness. Under these conditions, growth was completely suppressed, indicating that the organic carbon in the UF was insufficient to sustain biomass production. Therefore, the enhanced growth observed in UF-based cultures must be primarily attributed to photosynthetic activity and to the improved nutrient availability provided by the digestate-derived ultrafiltrate rather than to heterotrophic utilization of residual organic carbon. The ability of UF to promote biomass accumulation beyond that achieved with ammonium supplementation alone (Fig.1, Fig. 2, Fig. S1) suggests that digestate-derived ultrafiltrate acts as more than a simple nitrogen source. Indeed, UF provides a complex mixture of macro- and micronutrients, including potassium, phosphorus, iron and boron, which are absent or present at different concentrations in conventional synthetic media. The combined availability of these nutrients may improve cellular metabolism and photosynthetic performance, thereby supporting higher biomass productivity. Although the mechanisms responsible for this stimulatory effect remain to be elucidated, the data indicate that nutrient recovery through ultrafiltration generates a cultivation medium with properties that are potentially superior to those of standard synthetic formulations.

### 3.2 Ammonium and phosphate removal efficiency

Ammonium concentrations were measured at the beginning and end of the cultivation period to evaluate the nutrient-recovery capacity of *C. sorokiniana* and *N. oceanica* grown in ultrafiltrate (UF)-based media. Ammonium removal was quantified by distillation, and the results are reported in Table 2. Both microalgal species exhibited a high capacity to remove ammonium from the culture medium. *C. sorokiniana* achieved nearly complete ammonium removal in cultures containing 15% UF, whereas removal efficiency progressively declined as UF concentration increased. The lowest efficiency was observed in cultures supplemented with 42% UF, where approximately 64% of the initial ammonium was removed. Nevertheless, substantial ammonium depletion was observed across all treatments, indicating efficient nitrogen assimilation even under elevated nitrogen loads. Similarly, *N. oceanica* displayed remarkable ammonium-removal efficiency after 19 days of cultivation. At UF concentrations up to 15%, approximately 99% of the ammonium initially present in the medium was removed. Increasing UF concentration to 20% reduced removal efficiency to approximately 87%, although nitrogen recovery remained high. The decrease in removal efficiency observed at higher UF concentrations for both species is likely related to the increased ammonium availability, which exceeded the assimilation capacity associated with the observed growth rates.

**Table 2.** Ammonium removal efficiency of *Nannochloropsis oceanica* and *Chlorella sorokiniana* cultivated in media supplemented with increasing proportions of digestate-derived ultrafiltrate (UF). Removal efficiency was calculated from the difference between initial and final ammonium concentrations measured in the culture medium after 19 days of cultivation. Data are expressed as mean ± standard deviation (n = 3). Different letters within each species indicate significant differences among UF concentrations (one-way ANOVA followed by Tukey’s multiple-comparison test, p < 0.05).

| % UF | removal efficiency | % UF | removal efficiency |
| --- | --- | --- | --- |
| 5% | 98.31% ± 0.14<sup>a</sup> | 15% | 91% ± 0.98<sup>a</sup> |
| 10% | 99.40% ± 0.05<sup>a</sup> | 20% | 91% ± 0.95<sup>a</sup> |
| 15% | 99.63% ± 0.04<sup>a</sup> | 30% | 82% ± 1.44<sup>b</sup> |
| 20% | 87.46% ± 1.42<sup>b</sup> | 42% | 68% ± 0.54<sup>c</sup> |

To verify whether the recovered nitrogen was effectively incorporated into biomass, the elemental composition of the cells was analysed (Table S1). Total carbon content remained relatively constant in both species and cultivation conditions, ranging between 49 and 51% of dry weight. In *N. oceanica*, total nitrogen content was also largely unaffected by the growth medium (7.77 and 7.83% in BG-11 and UF-grown cells, respectively), suggesting that nitrogen recovered from the UF was assimilated with an efficiency comparable to that observed in synthetic media. In contrast, *C. sorokiniana* cultivated in UF exhibited a lower nitrogen content (6.35%) than cells grown in BG-11 (8.96%), despite producing substantially more biomass. This finding suggests that the enhanced biomass accumulation observed in UF-grown cultures was not accompanied by a proportional increase in cellular nitrogen content and may reflect changes in biomass composition.

The concentrations of macro- and micronutrients in the harvested biomass are reported in Table S2. Although some differences were observed between cells grown in synthetic media and UF, no major nutrient deficiencies were detected. In both species, phosphorus concentrations were lower in UF-grown biomass, whereas carbon content and growth performance remained largely unaffected, indicating efficient nutrient utilization. Likewise, reductions in the accumulation of certain micronutrients, including iron, copper, and zinc, did not negatively impact biomass productivity. Interestingly, species-specific responses were observed, such as the increased manganese accumulation in *C. sorokiniana* and the higher magnesium content in UF-grown *N. oceanica*. These results suggest that the distinct nutrient composition of UF affects cellular elemental profiles without compromising growth performance.

Overall, the high ammonium removal efficiencies achieved by both species, together with the maintenance of balanced elemental compositions and the substantial increase in biomass productivity, demonstrate that digestate-derived UF can effectively support both nutrient recovery and algal biomass production. These findings further support the use of UF as a sustainable alternative to conventional synthetic media within integrated nutrient-recovery and biomass-valorization processes.

### 3.3 Valorization of the biomass produced as biostimulant

The algal biomass produced using UF as nutrient source was then tested of its potential biostimulatory activity on plants, as a possible valorization strategy. Initially, a possible effect on germination was evaluated on tomato seeds (Fig. S4). Overall, treatment with microalgal species had a limited effect on tomato seed germination (Fig. S3). No consistent differences were observed between the treatment with microalgal biomass produced in BG-11 and biomass produced using UF. A decrease in germination percentage was observed at the highest concentration (25 g L⁻¹) for both microalgae species. Focusing on the effects on root length, *C. sorokiniana* biomass produced using UF showed a tendency to increase root length at concentrations ranging from 0.02 to 1 g L⁻¹, with the highest mean value recorded at 1 g L⁻¹ (Fig. S4). However, this value did not differ significantly from those of all corresponding controls. Biomass produced in BG-11 had only a limited effect on root length. For *N. oceanica*, higher mean root-length values were generally observed at concentrations ranging from 0.02 to 1 g L⁻¹ considering both growth conditions (Fig. S4), although the statistical significance of these differences depended on the treatment and reference control considered. At the highest concentrations, particularly at 25 g L⁻¹, root length tended to decrease for microalgal species (Fig. S4). Consistent with these findings previous works documented comparable dose-dependent effects on seed germination and early growth using *C. vulgaris*, *N. salina*, and *A. pratensis* in common bean (Elhakim et al., 2026). Several works reported the positive effects of the treatment with *C. sorokiniana* on the germination of seeds of different plant species as tomato and rice (Do et al., 2020). In lettuce, no effects on seed germination and dry biomass were observed in response to treatments with different concentrations of *C.* sorokiniana (Ribeiro et al., 2019). Less information is available regarding the biostimulant activity of *N. oceanica*.

A potential biostimulatory activity of microalgal biomass was then evaluated on hydroponically grown plants. Cultivation of tomato plants grown in hydroponic conditions, in which microalgal biomass was added to the nutrient solution, showed that *N. oceanica* positively affected root system development as well as root and shoot fresh weight (FW) in tomato seedlings. Similar effects were observed with biomass produced in either the synthetic medium (BG-11) or the ultrafiltrate (NUF) (Fig. S5). By contrast, the effects of *C. sorokiniana* varied more markedly depending on the growth medium, with biomass produced in BG-11 showing only limited effects on root-related parameters (Fig. S6). A previous work based on screening experiments on tomato plants demonstrated that microalgal effects are strongly species-dependent. The potential biostimulant activity of *C. sorokiniana* has already been reported in several studies (Martini et al., 2021; Mutale-Joan et al., 2020) including in relation to plant N nutrition (Chovancek et al., 2025). However, in the conditions herein tested, *C. sorokiniana* demonstrated a lower biostimulant effect compared to other species, specifically showing limited improvements in root length, shoot biomass, and nutrient accumulation, whereas extracts from the related marine species *N. gaditana* significantly enhanced all three parameters (Mutale-Joan et al., 2020).

We thus further evaluated the biostimulant potential of microalgal biomass, in relation to different N availability, focusing on *N. oceanica* produced using the UF. This species was selected because the initial hydroponic experiments (Fig. S5) indicated that biomass produced using the UF retained biostimulant activity comparable to that observed for biomass produced in the synthetic medium (BG-11). Furthermore, the potential application of *N. oceanica* as a plant biostimulant remains relatively underexplored.

To this end, hydroponic experiments were conducted in which tomato plants were grown in a full-strength nutrient solution (control) and a nutrient solution containing 30% of the standard N concentration (30% N). Under each nutritional condition, the effects of treatment with *N. oceanica* biomass produced using the ultrafiltrate on plant growth and root system development were evaluated. Compared with the corresponding untreated plants, NUF treatment increased several root parameters, particularly total root length and the number of secondary roots (i.e. root tips), irrespective of the N level (Fig. 4). Similar results were observed in hydroponically grown sugar beet treated with *Chlorella vulgaris* and *Scenedesmus quadricauda* extracts added to a complete nutrient solution nutrient solution (Barone et al., 2018). In addition, it was reported that applying a mixed microalgal consortium, composed mainly of *Chlorella* sp., *Scenedesmus* sp., *Spirulina* sp., and *Synechocystis* sp., to strawberry plants grown in vitro increased the number and length of their roots (Chaouch et al., 2023). A positive effect of *C. vulgaris* on tomato root initiation was reported when the microalgae species was cultivated both using a synthetic medium and treated sewage wastewater: the microalgae showed differences in auxin (IAA) and cytokinin (BA) levels in relation to the growth medium (Elakbawy et al., 2022). NUF treatment also increased shoot DW and both root FW and DW when plant grown under the standard N dose (100%) (Fig. 5). The treatment increases the DW of root also when the plant grown under a low N dose (30%) (Fig. 5). Although root parameters did not differ between plants grown at the full N dose and those grown at 30% of that dose, it is interesting that NUF treatment can increase the root-to-shoot ratio, both fresh and dry, particularly when at a lower N dose (Fig. 5). The positive effect on root biomass development relative to aboveground biomass is particularly important under stress conditions. Greater root system development can help explore a larger volume of soil, especially when nutrients are limited (Martini et al., 2021).

**Figure 4.**
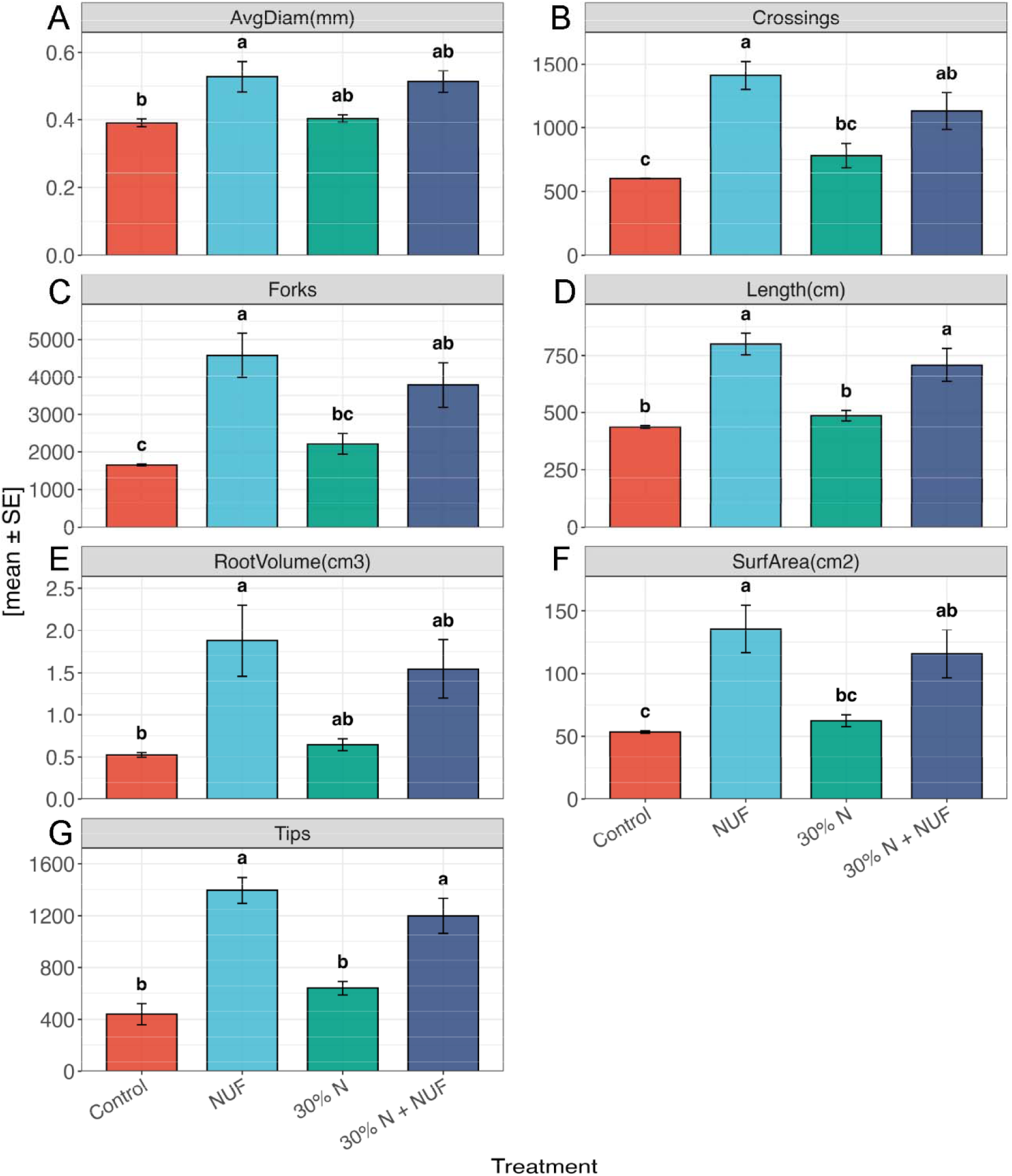
Root architectural responses of tomato plants to ultrafiltrate-derived *Nannochloropsis oceanica* biomass under contrasting nitrogen regimes. Average root diameter (A), number of crossings (B), number of forks (C), total root length (D), root volume (E), root surface area (F), and number of root tips (G) measured in tomato plants supplied with either full-strength nutrient solution (100% N) or reduced nitrogen availability (30% N), with or without treatment with *N. oceanica* biomass produced in digestate-derived ultrafiltrate (NUF). Data are expressed as mean ± standard error (n = 3). Different letters indicate statistically significant differences among treatments within each panel, according to one-way ANOVA followed by Tukey’s post-hoc test (p < 0.05).

**Figure 5.**
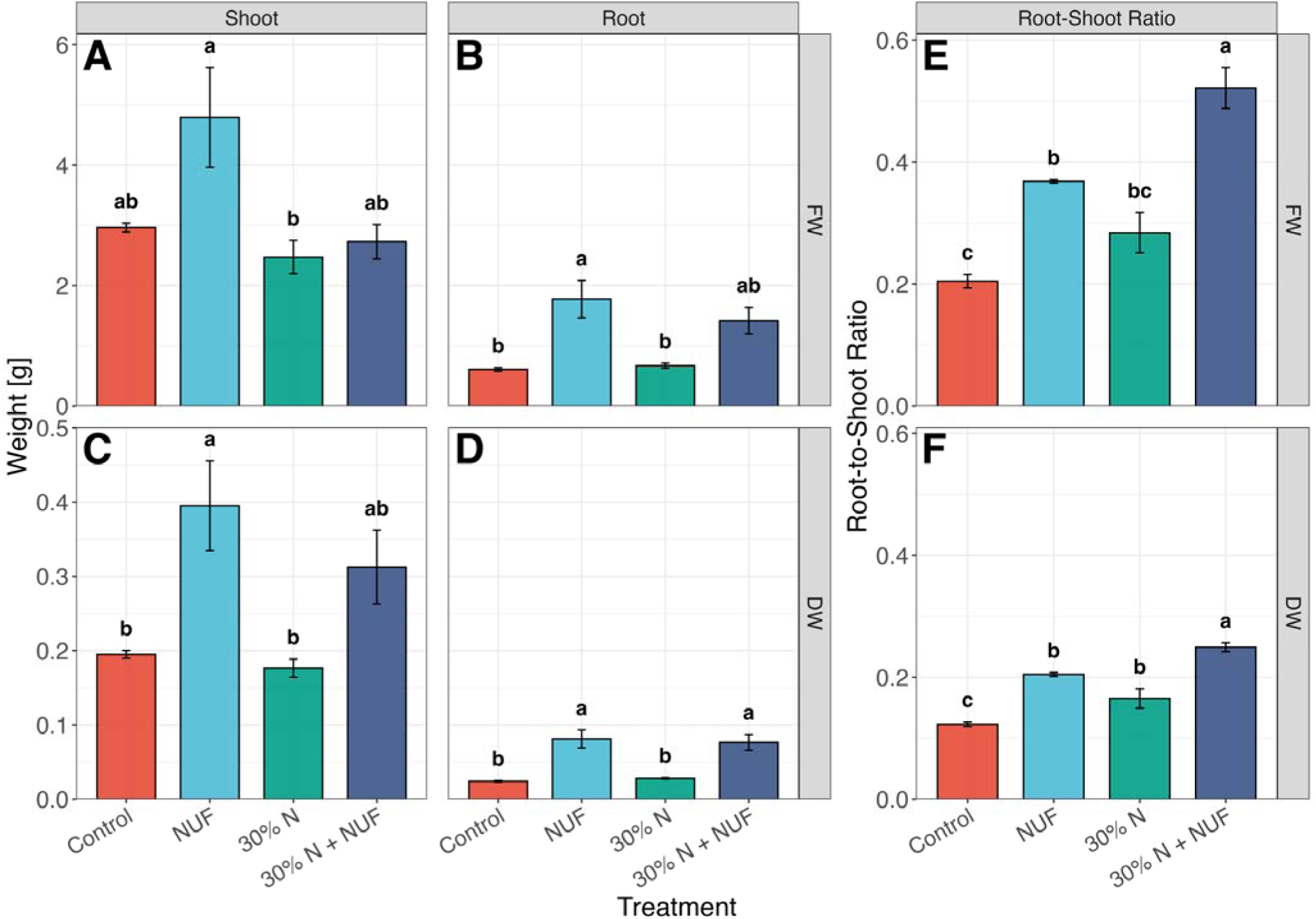
Growth-promoting effects of ultrafiltrate-derived *Nannochloropsis oceanica* biomass on tomato plants under contrasting nitrogen availability. Fresh shoot weight (A), fresh root weight (B), dry shoot weight (C), dry root weight (D), fresh root-to-shoot ratio (E), and dry root-to-shoot ratio (F) measured in tomato plants grown under full nitrogen supply (100% N) or reduced nitrogen availability (30% N), with or without treatment with *N. oceanica* biomass produced in digestate-derived ultrafiltrate (NUF). FW: fresh weight, DW: dry weight. Data are expressed as mean ± standard error (n = 3). Different letters indicate statistically significant differences among treatments within each panel, according to one-way ANOVA followed by Tukey’s post-hoc test (p < 0.05).

### 3.4 Towards a circular microalgae-based biorefinery for digestate valorization

The results obtained in this study demonstrate that livestock digestate ultrafiltrate should not be regarded merely as a waste-derived nutrient stream requiring treatment, but rather as a valuable resource capable of supporting the production of high-value microalgal biomass. Both *C. sorokiniana* and *N. oceanica* efficiently utilized the nutrients contained in the ultrafiltrate, achieving substantial ammonium removal while producing biomass levels significantly higher than those obtained in conventional synthetic media. These findings indicate that digestate-derived ultrafiltrate can simultaneously address two major challenges associated with anaerobic digestion systems: the management of nutrient-rich effluents and the economic sustainability of downstream valorization processes. Moreover, the twofold increase in biomass production using ultrafiltrate-derived media suggests this is a promising approach to optimizing algae cultivation, improving yield and reducing costs.

Importantly, the biomass generated during ultrafiltrate cultivation is not simply a by-product of the remediation process. Microalgal biomass represents a potential source of value-added agricultural biostimulants, creating an opportunity to reintroduce nutrients and bioactive compounds into crop production systems. From this perspective, the process investigated here establishes a closed-loop approach in which nutrients originating from livestock waste are recovered by microalgae, converted into biomass, and subsequently recycled back into agricultural production. Such integration increases resource-use efficiency, reduces dependence on synthetic fertilizers and cultivation media, and contributes to the development of circular bioeconomy strategies for the agricultural sector.

Particularly noteworthy is the ability of *C. sorokiniana* to maintain high productivity when ultrafiltrate was diluted only with water, without the addition of synthetic nutrients. This result suggests that the ultrafiltrate itself contains most of the components required to sustain microalgal growth and highlights the potential to drastically reduce nutrient input costs.

Combined with the high ammonium-removal efficiencies observed in both species, these findings support the feasibility of developing integrated systems in which digestate treatment, nutrient recovery, biomass production and biostimulant generation occur within a single process chain.

Overall, this study provides evidence that microalgae-based treatment of digestate ultrafiltrate can evolve from a wastewater remediation strategy into a multiproduct biorefinery platform capable of generating environmental and economic benefits simultaneously. Future work should focus on pilot-scale validation, comprehensive characterization of the bioactive compounds responsible for the biostimulant activity, and techno-economic assessments to evaluate the industrial implementation of this circular approach.

## 4. Conclusions

Livestock digestate ultrafiltrate proved to be an effective alternative to conventional synthetic media for the cultivation of both *Chlorella sorokiniana* and *Nannochloropsis oceanica*. The ultrafiltrate supported robust growth, high biomass productivity, and efficient ammonium removal, demonstrating its suitability as a nutrient-rich substrate for microalgal cultivation. Biomass production increased by up to 133% in *C. sorokiniana* and 104% in *N. oceanica* compared with the respective synthetic controls, while ammonium removal exceeded 98% under the most favorable conditions. Furthermore, *C. sorokiniana* successfully grew in ultrafiltrate diluted only with water, indicating that the ultrafiltrate can provide most of the nutrients required for cultivation without the addition of synthetic media components.

These results demonstrate that ultrafiltration-derived digestate fractions can simultaneously support nutrient recovery and enhanced microalgal biomass production. The biomass generated through this process represents a promising feedstock for biostimulant applications, further increasing the value of digestate-derived resources. Overall, this work identifies digestate ultrafiltrate as a low-cost cultivation medium with the potential to improve the sustainability and economic feasibility of microalgal production systems. Importantly, the enhanced biomass productivity observed in UF-grown cultures cannot be explained solely by nitrogen availability, suggesting that the complex nutrient composition of the digestate-derived ultrafiltrate contributes to the stimulation of microalgal growth.

## Data Availability Statement

All data and results are presented in the paper and in the Supporting Information.

## Acknowledgements

Authors thank the Centro Piattaforme Tecnologiche for providing access to the core facilities of the University of Verona.

## Bibliography

Alice, F., Lusine, M., Sofia, C., Belina, R., Daniel, F., Gayane, A., Luisa, G. 2021. Biostimulant and biopesticide potential of microalgae growing in piggery wastewater. Environmental Advances, 4(100062).

Allen, M.M., Stanier, R.Y. 1968. Growth and division of some unicellular blue-green algae. J Gen Microbiol, 51(2), 199–202.

Barone, V., Baglieri, A., Stevanato, P., Broccanello, C., Bertoldo, G., Bertaggia, M., Cagnin, M., Pizzeghello, D., Moliterni, V., Mandolino, G., Fornasier, F., Squartini, A., Nardi, S., Concheri, G. 2018. Root morphological and molecular responses induced by microalgae extracts in sugar beet (*Beta vulgaris* L.). Journal of Applied Phycology, 30(2), 1061–1071.

Bauer, L., Ranglová, K., Masojídek, J., Drosg, B., Meixner, K. 2021. Digestate as Sustainable Nutrient Source for Microalgae—Challenges and Prospects. Applied Sciences, 11(3), 1056.

Bernhard, D., Werner, F., Teodorita, A.S., Michael, M., Bernd, L. 2015. Nutrient Recovery by Biogas Digestate Processing, (Ed.) I. Bioenergy, pp. 39.

Brooke, F., Ma, Q., Rahimi, M., Silkina, A., Constantin, C., Badulescu, L., La Terza, A., Turkmen, B., Bajoub, A., Ajal, E., van Mullem, D., Halim, R. 2025. Nannochloropsis for the bioremediation of brewery side streams and co-production of aquaculture feed and bio-Fertiliser: A comprehensive review. Algal Research-Biomass Biofuels and Bioproducts, 89.

Cai, T., Ge, X., Park, S., Li, Y. 2013. Comparison of *Synechocystis* sp PCC6803 and *Nannochloropsis salina* for lipid production using artificial seawater and nutrients from anaerobic digestion effluent. Bioresource Technology, 144, 255–260.

Canini, D., Martini, F., Cazzaniga, S., Miotti, T., Pacenza, B., D’Adamo, S., Ballottari, M. 2024. Genetic engineering of Nannochloropsis oceanica to produce canthaxanthin and ketocarotenoids. Microb Cell Fact, 23(1), 322.

Cavinato, C., Fatone, F., Bolzonella, D., Pavan, P. 2010. Thermophilic anaerobic co-digestion of cattle manure with agro-wastes and energy crops: Comparison of pilot and full scale experiences. Bioresource Technology, 101(2), 545–550.

Chaouch, R., Kthiri, Z., Soufi, S., Jabeur, M.B., Bettaieb, T.a. 2023. Assessing the biostimulant effect of micro-algae and thyme essential oil during in-vitro and ex-vitro rooting of strawberry. South African Journal of Botany, 162, 9.

Chiaiese, P., Corrado, G., Colla, G., Kyriacou, M., Rouphael, Y. 2018. Renewable Sources of Plant Biostimulation: Microalgae as a Sustainable Means to Improve Crop Performance. Frontiers in Plant Science, 9.

Chojnacka, K., Chojnacki, M. 2024. Nutrient recovery from anaerobic digestate: Fertilizer informatics for circular economy. Environmental Research, 245.

Chovancek, E., Poque, S., Bayram, E., Borhan, E., Jokel, M., Rantanen, I., Haznedaroglu, B., Himanen, K., Sirin, S., Allahverdiyeva, Y. 2025. Stepwise processing of *Chlorella sorokiniana* confers plant biostimulant that reduces mineral fertilizer requirements. Bioresource Technology, 418.

Chuka-ogwude, D., Ogbonna, J., Moheimani, N. 2020. A review on microalgal culture to treat anaerobic digestate food waste effluent. Algal Research-Biomass Biofuels and Bioproducts, 47.

Collos, Y., Harrison, P. 2014. Acclimation and toxicity of high ammonium concentrations to unicellular algae. Marine Pollution Bulletin, 80(1-2), 8–23.

Dahmani, S., Zerrouki, D., Ramanna, L., Rawat, I., Bux, F. 2016. Cultivation of Chlorella pyrenoidosa in outdoor open raceway pond using domestic wastewater as medium in arid desert region. Bioresource Technology, 219, 749–752.

De-Bashan, L., Trejo, A., Huss, V., Hernandez, J., Bashan, Y. 2008. *Chlorella sorokiniana* UTEX 2805, a heat and intense, sunlight-tolerant microalga with potential for removing ammonium from wastewater. Bioresource Technology, 99(11), 4980–4989.

Do, T., Tran, D., Le, T., Nguyen, Q. 2020. Characterization of Endogenous Auxins and Gibberellins Produced by *Chlorella sorokiniana* TH01 under Phototrophic and Mixtrophic Cultivation Modes toward Applications in Microalgal Biorefinery and Crop Research. Journal of Chemistry, 2020.

Elakbawy, W.M., Shanab, S.M.M., Shalaby, E.A. 2022. Enhancement of plant growth regulators production from microalgae cultivated in treated sewage wastewater (TSW). BMC Plant Biol, 22(1), 377.

Elhakim, K., Gharib, F., Ahmed, E. 2026. Agronomic evaluation of microalgal extracts as seed bio stimulants for improving germination and early growth of common bean. Scientific Reports, 16(1).

Fernandes, F., Silkina, A., Fuentes-Grünewald, C., Wood, E.E., Ndovela, V.L.S., Oatley-Radcliffe, D.L., Lovitt, R.W., Llewellyn, C.A. 2020. Valorising nutrient-rich digestate: Dilution, settlement and membrane filtration processing for optimisation as a waste-based media for microalgal cultivation. Waste Manag, 118, 197–208.

Freudenberg, R., Baier, T., Einhaus, A., Wobbe, L., Kruse, O. 2021. High cell density cultivation enables efficient and sustainable recombinant polyamine production in the microalga Chlamydomonas reinhardtii. Bioresource Technology, 323.

Fuentes-Grünewald, C., Gayo-Peláez, J., Ndovela, V., Wood, E., Kapoore, R., Llewellyn, C. 2021. Towards a circular economy: A novel microalgal two-step growth approach to treat excess nutrients from digestate and to produce biomass for animal feed. Bioresource Technology, 320.

Giuliano, A., Bolzonella, D., Pavan, P., Cavinato, C., Cecchi, F. 2013. Co-digestion of livestock effluents, energy crops and agro-waste: Feeding and process optimization in mesophilic and thermophilic conditions. Bioresource Technology, 128, 612–618.

Gu, Z., Liu, Y., Zou, G., Zhang, Q., Lu, R., Yan, H., Cao, L., Liu, T., Ruan, R. 2021. Enhancement of nutrients removal and biomass accumulation of Chlorella vulgaris in pig manure anaerobic digestate effluent by the pretreatment of indigenous bacteria. Bioresour Technol, 328, 124846.

Gupta, S., Pawar, S., Pandey, R. 2019. Current practices and challenges in using microalgae for treatment of nutrient rich wastewater from agro-based industries. Science of the Total Environment, 687, 1107–1126.

Hairuddin, N., Mohamed, R., Arifin, S., Adrin, M., Azmi, N. 2025. Recent Trends in the Dual Role of Microalgae in Wastewater Remediation and Bioproduct Generation for Aquafeed. Journal of Environmental Science and Management, 28(2).

Herbes, C., Roth, U., Wulf, S., Dahlin, J. 2020. Economic assessment of different biogas digestate processing technologies: A scenario-based analysis. Journal of Cleaner Production, 255.

Huete-Ortega, M., Okurowska, K., Kapoore, R., Johnson, M., Gilmour, D., Vaidyanathan, S. 2018. Effect of ammonium and high light intensity on the accumulation of lipids in *Nannochloropsis oceanica* (CCAP 849/10) and *Phaeodactylum tricornutum* (CCAP 1055/1). Biotechnology For Biofuels, 11.

Kapoore, R., Wood, E., Llewellyn, C. 2021. Algae biostimulants: A critical look at microalgal biostimulants for sustainable agricultural practices. Biotechnology Advances, 49.

Leong, Y., Chang, J. 2022. Integrated role of algae in the closed-loop circular economy of anaerobic digestion. Bioresource Technology, 360.

Li, Y., Chen, Y., Chen, P., Min, M., Zhou, W., Martinez, B., Zhu, J., Ruan, R. 2011. Characterization of a microalga *Chlorella* sp well adapted to highly concentrated municipal wastewater for nutrient removal and biodiesel production. Bioresource Technology, 102(8), 5138–5144.

Liang, W., Min, M., Yecong, L., Paul, C., Yifen, C., Yuhuan, L., Yingkuan, W., Roger, R. 2010. Cultivation of Green Algae C*hlorella* sp. in Different Wastewaters from Municipal Wastewater Treatment Plant. Appl Biochem Biotechnol, 162, 1174–1186.

Ljumovic, K., Betterle, N., Baietta, A., Ballottari, M. 2024. Valorization of wastewater from industrial hydroponic cultivations using the microalgal species *Chlorella vulgaris*. Algal Research-Biomass Biofuels and Bioproducts, 81.

Mancuso, G., Habchi, S., Maraldi, M., Valenti, F., El Bari, H. 2024. Comprehensive review of technologies for separate digestate treatment and agricultural valorisation within circular and green economy. Bioresource Technology, 409.

Manzo, R., Papirio, S., Esposito, G., Matassa, S. 2026. Enabling protein-rich microalgae production from anaerobic digestate through direct aeration stripping for gaseous nitrogen and carbon supply. Waste Manag, 209, 115187.

Markou, G., Muylaert, K. 2016. Effect of light intensity on the degree of ammonia toxicity on PSII activity of *Arthrospira platensis* and *Chlorella vulgaris*. Bioresource Technology, 216, 453–461.

Martini, F., Beghini, G., Zanin, L., Varanini, Z., Zamboni, A., Ballottari, M. 2021. The potential use of Chlamydomonas reinhardtii and Chlorella sorokiniana as biostimulants on maize plants. Algal Res, 60.

Miotti, T., Perozeni, F., Peli, M., Bellamoli, F., Lo Coco, R., Frison, N., Zamboni, A., Felis, G., Ballottari, M. 2026. Valorization of anaerobic digestate as nutrients source for *Chlorella vulgaris* to produce biostimulants: Analysis of photosynthetic response and microbiome composition. Algal Research-Biomass Biofuels and Bioproducts, 96.

Morillas-España, A., Ruiz-Nieto, Á., Lafarga, T., Acién, G., Arbib, Z., González-López, C.V. 2022. Biostimulant Capacity of Chlorella and Chlamydopodium Species Produced Using Wastewater and Centrate. Biology (Basel), 11(7).

Mutale-Joan, C., Redouane, B., Najib, E., Yassine, K., Lyamlouli, K., Laila, S., Zeroual, Y., Hicham, E.A. 2020. Screening of microalgae liquid extracts for their bio stimulant properties on plant growth, nutrient uptake and metabolite profile of Solanum lycopersicum L. Sci Rep, 10(1), 2820.

Pizzera, A., Scaglione, D., Bellucci, M., Marazzi, F., Mezzanotte, V., Parati, K., Ficara, E. 2019. Digestate treatment with algae-bacteria consortia: A field pilot-scale experimentation in a sub-optimal climate area. Bioresource Technology, 274, 232–243.

Praveen, P., Guo, Y., Kang, H., Lefebvre, C., Loh, K. 2018. Enhancing microalgae cultivation in anaerobic digestate through nitrification. Chemical Engineering Journal, 354, 905–912.

Ramasamy, S., Pakshirajan, K., Murugan, D., Saini, G.K. 2024. Lutein Production by Halophilic Microalgae Using Anaerobic Digestate as the Substrate and Its Potential Application as a Biopesticide. Appl Biochem Biotechnol, 196(5), 2591–2611.

Ribeiro, D.M., Zanetti, G.T., Julião, M.H.M., Masetto, T.E., Gelinski, J.M.L.N., Fonseca, G.G. 2019. Effect of different culture media on growth of Chlorella sorokiniana and the influence of microalgal effluents on the germination of lettuce seed. Journal of Applied Biology & Biotechnology, 7(1), 5.

Rizzioli, F., Bertasini, D., Bolzonella, D., Frison, N., Battista, F. 2023. A critical review on the techno-economic feasibility of nutrients recovery from anaerobic digestate in the agricultural sector. Separation and Purification Technology, 306.

Rizzioli, F., Cirilli, M., Frison, N., Bolzonella, D., Battista, F. 2025. Nutrient recovery from anaerobic digestate by different combination of pressure driven membranes. Journal of Cleaner Production, 494.

Rossi, S., Mantovani, M., Marazzi, F., Mezzanotte, V., Ficara, E. 2025. Long-term outdoor operation of microalgae-based digestate treatment: impact of external drivers on process performances and techno-economic assessment. Bioresource Technology, 427.

Sanz-Luque, E., Chamizo-Ampudia, A., Llamas, A., Galvan, A., Fernandez, E. 2015. Understanding nitrate assimilation and its regulation in microalgae. Front Plant Sci, 6, 899.

Sasongko, N.A., Ayre, J., Noguchi, R., Moheimani, N.R., Bahri, P.A., Anda, M., Nakajima, M., Soekotjo, E.S.A., Pertiwi, G.A., Handayani, T., Steven, S. 2025. Utilization and integration of microalgae consortium in treating undiluted anaerobic digestate animal effluent to produce animal feed, bio-oil, and biogas. Bioresour Technol, 416, 131788.

Scarponi, P., Ghirardini, A., Bravi, M., Cavinato, C. 2021. Evaluation of *Chlorella vulgaris* and *Scenedesmus obliquus* growth on pretreated organic solid waste digestate. Waste Management, 119, 235–241.

Sharma, R., Choudhary, P., Thakur, G., Pathak, A., Singh, S., Kumar, A., Lo, S., Kumar, P. 2025. Sustainable management of biowaste to bioenergy: A critical review on biogas production and techno-economic challenges. Biomass & Bioenergy, 196.

Sobolewska, E., Borowski, S., Nowicka-Krawczyk, P., Jurczak, T. 2023. Growth of microalgae and cyanobacteria consortium in a photobioreactor treating liquid anaerobic digestate from vegetable waste. Sci Rep, 13(1), 22651.

Stiles, W., Styles, D., Chapman, S., Esteves, S., Bywater, A., Melville, L., Silkina, A., Lupatsch, I., Grünewald, C., Lovitt, R., Chaloner, T., Bull, A., Morris, C., Llewellyn, C. 2018. Using microalgae in the circular economy to valorise anaerobic digestate: challenges and opportunities. Bioresource Technology, 267, 732–742.

Tawfik, A., Eraky, M., Alhajeri, N., Osman, A., Rooney, D. 2022. Cultivation of microalgae on liquid anaerobic digestate for depollution, biofuels and cosmetics: a review. Environmental Chemistry Letters, 20(6), 3631–3656.

Terrado, R., Monier, A., Edgar, R., Lovejoy, C. 2015. Diversity of nitrogen assimilation pathways among microbial photosynthetic eukaryotes. J Phycol, 51(3), 490–506.

Uggetti, E., Sialve, B., Latrille, E., Steyer, J. 2014. Anaerobic digestate as substrate for microalgae culture: The role of ammonium concentration on the microalgae productivity. Bioresource Technology, 152, 437–443.

Veronesi, D., Idà, A., D’Imporzano, G., Adani, F., Pierucci, S., Klemes, J. 2015. Microalgae Cultivation: Nutrient Recovery from Digestate for Producing Algae Biomass. Icheap12: 12th International Conference on Chemical & Process Engineering, 43, 1201–1206.

Wester-Larsen, L., Müller-Stöver, D., Salo, T., Jensen, L. 2022. Potential ammonia volatilization from 39 different novel biobased fertilizers on the European market - A laboratory study using 5 European soils. Journal of Environmental Management, 323.

Xia, A., Murphy, J. 2016. Microalgal Cultivation in Treating Liquid Digestate from Biogas Systems. Trends in Biotechnology, 34(4), 264–275.

Yan, F. 2025. Agricultural biomass wastes and their resource utilization technologies: A review. Biomass & Bioenergy, 203.

Zhu, S., Jiang, R., Qin, L., Huang, D., Yao, C., Xu, J., Wang, Z. 2022. Integrated strategies for robust growth of *Chlorella vulgaris* on undiluted dairy farm liquid digestate and pollutant removal. Science of the Total Environment, 852.

Álvarez-González, A., Serrano, L., Gorchs, G., Uggetti, E. 2025. Exploring the biostimulant potential of Scenedesmus sp. grown in wastewater: impacts on plant growth and photosynthetic activity of lettuce. Chemosphere, 382, 144494.

