## Supporting Information for "Agricultural digestate ultrafiltrate outperforms synthetic media for microalgal biomass production while enabling biostimulant generation"

The following Supporting Information is available for this article:

Table S1. Carbon and nitrogen content of C. sorokiniana and N. oceanica biomass grown in BG-11 and UF media.

Table S2. Mineral nutrient composition of microalgal biomass cultivated in synthetic and ultrafiltrate-based media

Fig. S1. *Chlorella sorokiniana* and *Nannochloropsis oceanica* tolerance to $NH_4HCO_3$.

Fig. S2. *Nannochloropsis oceanica* dependence on ASW salts in the ultrafiltrate.

Fig. S3. Identification of mineral nutrients limiting *Nannochloropsis oceanica* growth in ultrafiltrate-based media.

Fig. S4. Biostimulant activity of *Chlorella sorokiniana* and *Nannochloropsis oceanica* biomass on tomato seed germination and root growth.

Fig. S5. Hydroponic treatment of tomato seedlings with *Nannochloropsis oceanica*

Fig. S6. Hydroponic treatment of tomato seedlings with *Chlorella sorokiniana*

**Table S1. Carbon and nitrogen content of C. sorokiniana and N. oceanica biomass grown in BG-11 and UF media.** Total N and C determined in *C. sorokiniana* and *N. oceanica* grown using synthetic medium (BG-11) and ultrafiltrate (UF). Data are expressed as mean ± standard deviation (n = 3).

| | **Total N (%)** | **Total C (%)** |
|---|---|---|
| *C. sorokiniana* (BG-11) | 8.96 ± 0.64 | 50.83 ± 2.80 |
| *C. sorokiniana* (UF) | 6.35 ± 0.99 | 49.20 ± 0.35 |
| *N. oceanica* (BG-11) | 7.77 ± 0.08 | 49.83 ± 0.46 |
| *N. oceanica* (UF) | 7.83 ± 0.75 | 50.57 ± 2.91 |

**Table S2. Mineral nutrient composition of microalgal biomass cultivated in synthetic and ultrafiltrate-based media.** Macro- and micronutrient concentrations of *C. sorokiniana* and *N. oceanica* grown using synthetic medium (BG-11) and ultrafiltrate (UF). Data are expressed as mean ± standard deviation (n = 3).

| | ***C. sorokiniana* (BG-11)** | ***C. sorokiniana* (UF)** | ***N. oceanica* (BG-11)** | ***N. oceanica* (UF)** |
|---|---|---|---|---|
| Ca (ppm) | 646.59 ± 111.96 | 216.09 ± 77.14 | 401.37 ± 31.63 | 477.63 ± 266.62 |
| K (ppm) | 16139.24 ± 308.32 | 7212.99 ± 87.35 | 15138.79 ± 438.87 | 16206.66 ± 2959.40 |
| Mg (ppm) | 2653.74 ± 86.58 | 1197.67 ± 46.81 | 3353.99 ± 70.44 | 4002.05 ± 820.72 |
| P (ppm) | 8636.19 ± 124.26 | 4941.36 ± 78.46 | 9888.33 ± 210.77 | 4448.39 ± 887.58 |
| B (ppm) | 2.34 ± 0.26 | n.d. | n.d. | 6.29 ± 2.68 |
| Cu (ppm) | 25.05 ± 0.35 | 4.53 ± 0.11 | 6.52 ± 0.09 | 2.61 ± 0.83 |
| Fe (ppm) | 574.27 ± 6.21 | 175.95 ± 2.46 | 1493.50 ± 27.57 | 212.09 ± 38.31 |
| Mn (ppm) | 94.06 ± 0.98 | 113.31 ± 2.20 | 167.36 ± 2.93 | 40.69 ± 7.54 |
| Mo (ppm) | 0.63 ± 0.01 | 0.45 ± 0.01 | 0.44 ± 0.01 | 0.40 ± 0.07 |
| Ni (ppm) | 0.79 ± 0.35 | 2.23 ± 0.06 | 0.28 ± 0.18 | 0.65 ± 0.29 |
| Zn (ppm) | 72.65 ± 4.47 | 25.16 ± 0.93 | 161.41 ± 2.37 | 59.42 ± 12.23 |

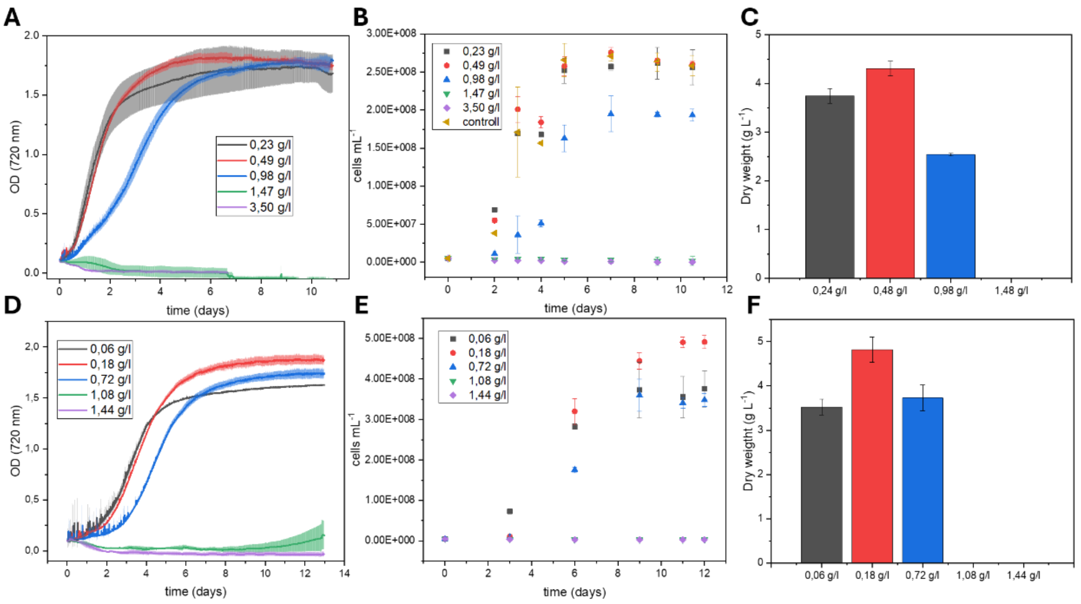


**Fig. S1. *Chlorella sorokiniana* and *Nannochloropsis oceanica* tolerance to $NH_4HCO_3$.** *C. sorokiniana* (A-C) and *N. oceanica* (D-F) were grown at different concentrations of $NH_4HCO_3$ at 24° C, bubbling 3% $CO_2$ with 1000 µmol photons m$^{-2}$ s$^{-1}$ of white light in BG11 and ASW medium for *C. sorokiniana* and *N. oceanica*, respectively. The black curves correspond to the concentration of nitrogen used in BG11 and ASW; the concentration is indicated as g/l of nitrogen. The growth was monitored by measuring optical density at 720 nm (A, D) and cell counts (B, D). On the last day of the experiment, the dry weight was determined (C, F). Error bars are reported as standard deviations (n=2).

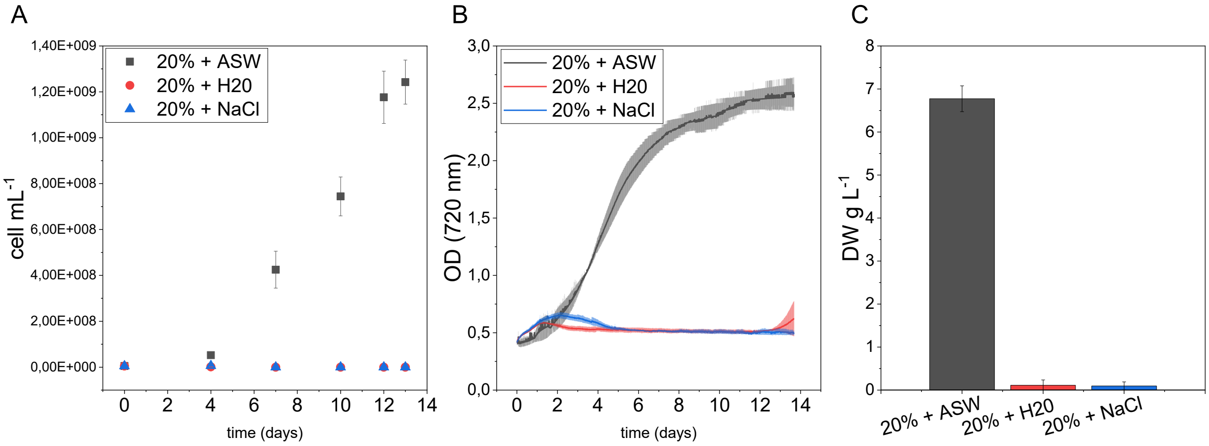


**Fig. S2. *Nannochloropsis oceanica* dependence on ASW salts in the ultrafiltrate.** (A) *N. oceanica* grown at 20% of UF in ASW medium at 24° C, bubbling 3% CO2, adding ASW salts (black), diluting UF in water (red) or adding NaCl (24 g/l). The growth was monitored by measuring optical density at 720 nm (A) and cell count (B). On the last day, the dry weight was determined (C). Error bars are reported as standard deviations (n=2).

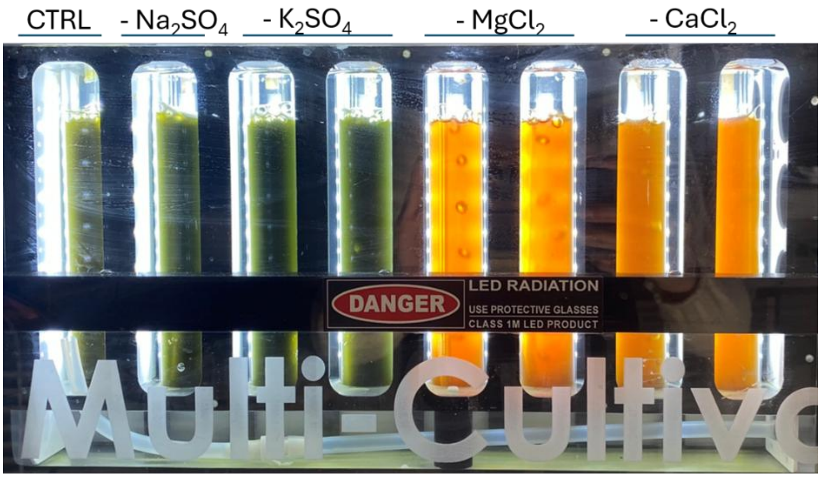


**Fig. S3. Identification of mineral nutrients limiting *Nannochloropsis oceanica* growth in ultrafiltrate-based media.** Representative cultures of *N. oceanica* grown in 20% ultrafiltrate (UF) diluted with artificial seawater (ASW) and in media where individual ASW salts were sequentially omitted and replaced by UF. From left to right: complete ASW + 20% UF (CTRL), ASW without $Na_2SO_4$ + 20% UF, ASW without $K_2SO_4$ + 20% UF, ASW without $MgCl_2$ + 20% UF, and ASW without $CaCl_2$ + 20% UF. Cultures lacking $MgCl_2$ or $CaCl_2$ exhibited severe growth inhibition, as indicated by the absence of the characteristic green pigmentation and the persistence of the orange color of the ultrafiltrate. In contrast, omission of $Na_2SO_4$ or $K_2SO_4$ had only minor effects on biomass accumulation. These results indicate that magnesium and calcium supplied by the artificial seawater medium are essential for *N. oceanica* growth and cannot be provided in sufficient amounts by the digestate-derived ultrafiltrate alone.

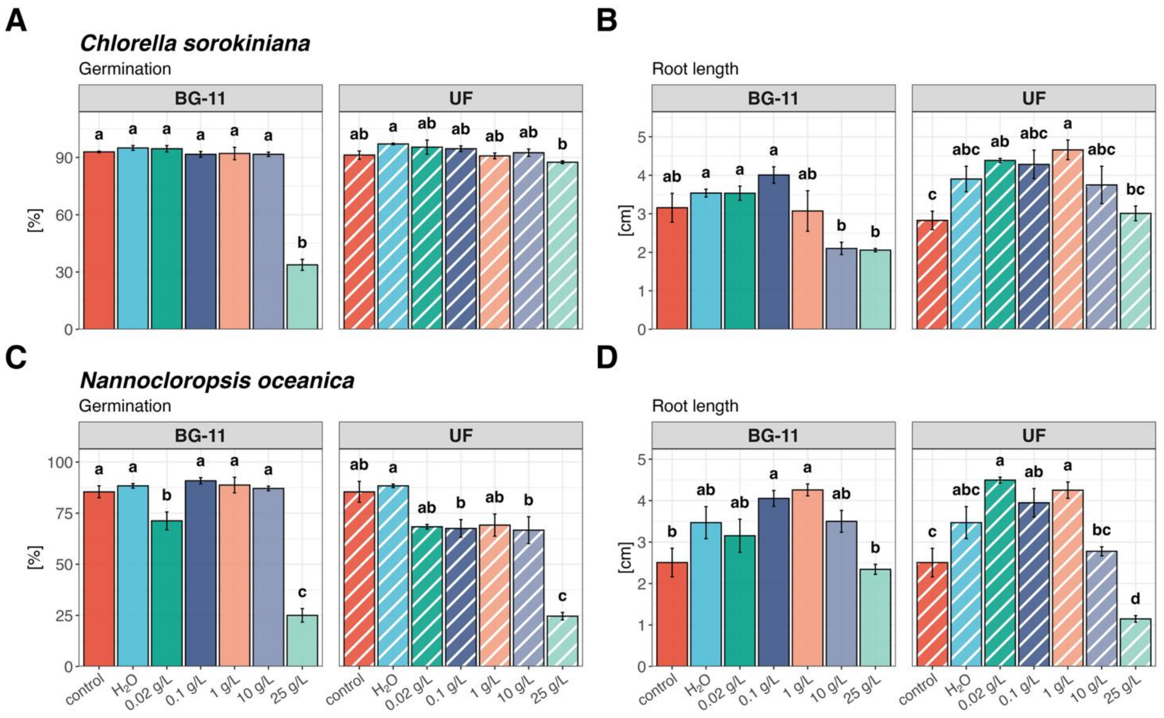


**Fig. S4. Biostimulant activity of *Chlorella sorokiniana* and *Nannochloropsis oceanica* biomass on tomato seed germination and root growth.** Germination percentage and root length of untreated tomato seeds (control), primed with $H_2O$, or primed with solutions containing increasing concentrations (0.02, 0.1, 1, 10, and 25 g $L^{-1}$) of *C. sorokiniana* or *N. oceanica* biomass, grown either in synthetic medium (BG-11) or using the ultrafiltrate (UF). (A) Germination (%) and (B) root length (cm) of seedlings primed with *C. sorokiniana*; (C) germination (%) and (D) root length (cm) of seedlings primed with *N. oceanica*. Data are expressed as mean ± standard error (n = 3); different letters indicate statistically significant differences among treatments within each panel, as determined by one-way ANOVA followed by Tukey's post-hoc test ($p < 0.05$).

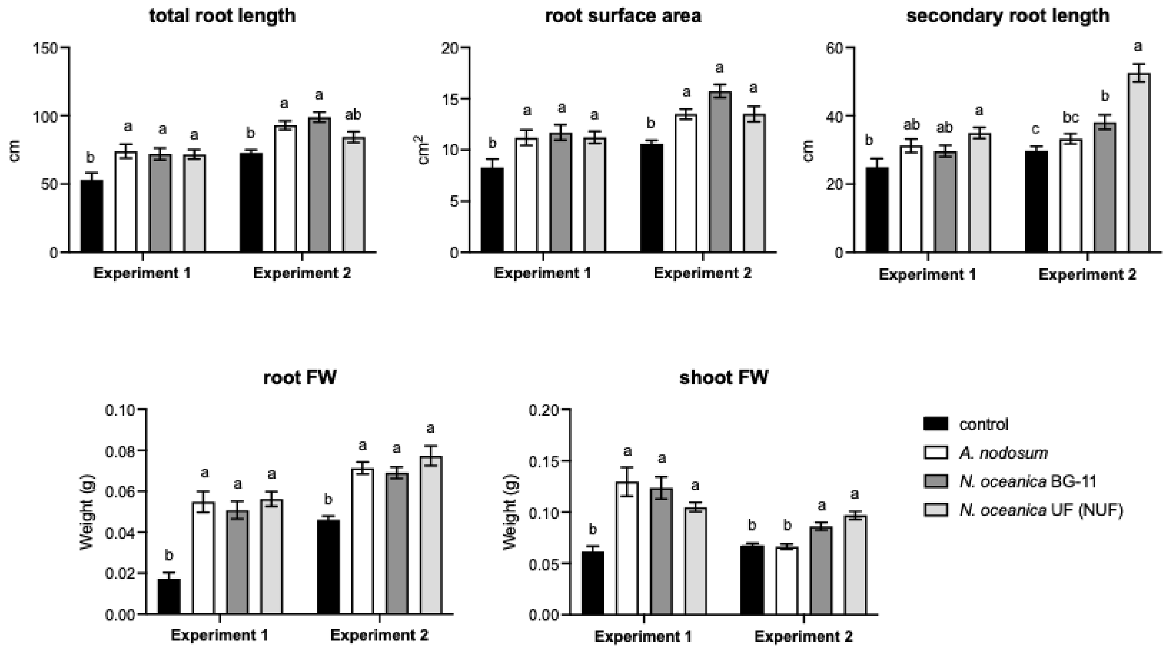


**Fig. S5. Hydroponic treatment of tomato seedlings with *Nannochloropsis oceanica.*** Total root length, root surface area, secondary root length, root FW and shoot FW of untreated tomato seedlings (control), seedlings treated with *A. nodosum*, and seedlings treated with *N. oceanica* grown in synthetic medium (BG-11) and using the ultrafiltrate (NUF). Data are expressed as mean ± standard error (n = 40); different letters indicate statistically significant differences among treatments within each panel, as determined by one-way ANOVA followed by Tukey's post-hoc test ($p < 0.05$).

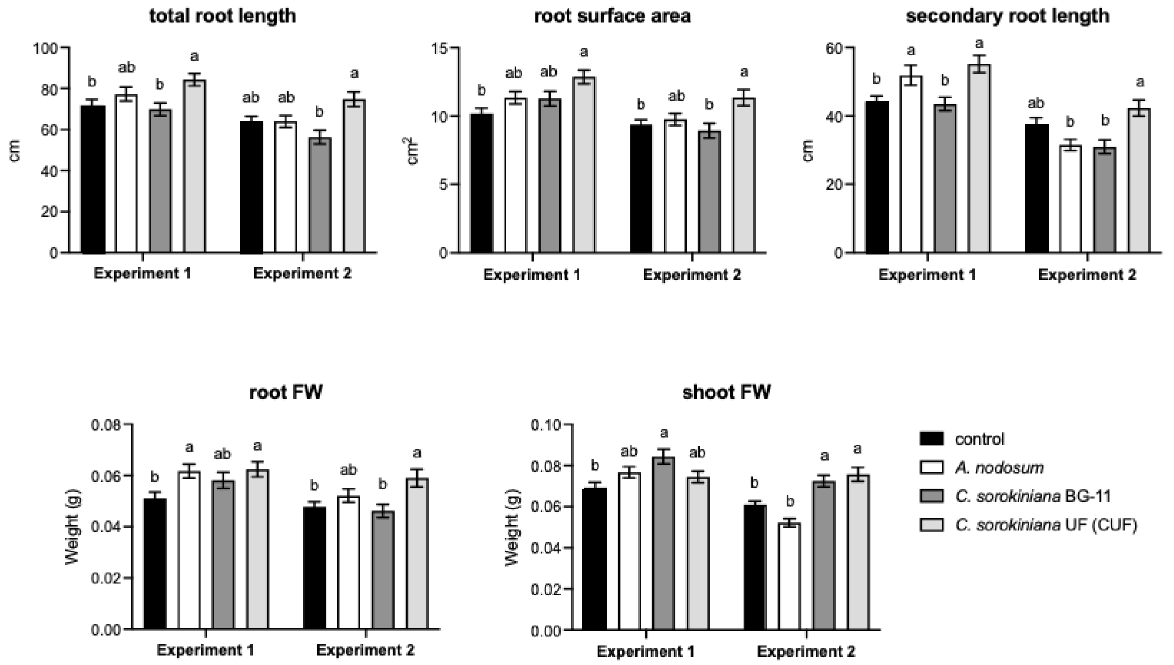


**Fig. S6. Hydroponic treatment of tomato seedlings with *Chlorella sorokiniana.*** Total root length, root surface area, secondary root length, root FW and shoot FW of untreated tomato seedlings (control), seedlings treated with *A. nodosum*, and seedlings treated with *C. sorokiniana* grown in synthetic medium (BG-11) and using the ultrafiltrate (CUF). Data are expressed as mean ± standard error (n = 40); different letters indicate statistically significant differences among treatments within each panel, as determined by one-way ANOVA followed by Tukey's post-hoc test ($p < 0.05$).
